# Transcriptomic data and biomedical literature synergize in finding pharmacologic gene regulators

**DOI:** 10.64898/2026.03.13.708862

**Authors:** Cole A. Deisseroth, Bess Brazelton, Zahid Shaik, Benjamin S. Sandberg, Zhandong Liu, Huda Y. Zoghbi

**Author notes:** Corresponding authors: Huda Y. Zoghbi,; Zhandong Liu.

## Abstract

Most disorders caused by a deficiency or excess of one gene product lack targeted therapies. Since these disorders can be modeled with a gene overexpression, knockout, or knockdown, drugs that oppose the transcriptomic effects of such perturbations may be promising therapeutic candidates. RNA-Sequencing (RNA-Seq) studies can fuel this drug-prioritization, but their labels, written in plain language, must be annotated manually. Hence, we introduce Signature-based Networks from Automatically Curated Knockout, Knockdown, and Small-molecule Studies (SNACKKSS), which automatically curates gene-disruption and drug-treatment studies from the Gene Expression Omnibus and, in partnership with uniformly computed read count datasets, feeds the labels and RNA-Seq data directly into regulatory relationship predictions. Through cross-validation, we show that SNACKKSS’ predictions (specifically, from a variant called “SA4”) make a unique contribution to finding protein-inhibiting compounds, even alongside existing predictors. We demonstrate the benefit of aggregating multiple predictive tools, and provide this powerful ensemble alongside SNACKKSS. Importantly, we advise researchers to test complex machine learning models on multiple devices, since the hardware architecture can affect their output. Nonetheless, our findings support the value of integrating automatically curated RNA-Seq signatures with literature- and co-expression-based predictors for drug-repurposing prioritization.

## Introduction

Many human diseases are caused by insufficient or excessive activity from specific proteins. Examples include cancer (hyperactive oncogenes and deficient tumor suppressors), endocrine disorders (releasing too much or too little of a given hormone, or failing to properly respond to it), and Mendelian diseases (hypo- or hypermorphic variants in one gene). Mendelian disorders are particularly challenging to treat, due to the sheer number of potential protein defects, and the vast majority of these proteins not having a known safe treatment that can support or inhibit them. One would ideally target the protein directly, as opposed to simply managing symptoms. Targeted gene therapies (ongoing efforts reviewed by Laurent et al.(1)) and engineered molecules that specifically address the defect(2, 3) are in active development, with some now having Food and Drug Administration (FDA) approval(4). Another approach is drug repurposing—finding chemicals with well-established toxicity profiles, originally intended for a different disorder, that also happen to regulate the affected gene product. General examples are reviewed by Sun et al.(5), and Dalton et al.(6) and Hope et al.(7) describe repurposing screens in fruit flies to find treatments for congenital disorders of glycosylation. If there is a molecule with a good safety profile that is particularly likely to address the deficiency or excess of a given gene product, it could be tested and brought to the bedside quickly(5). Therefore, a bioinformatic tool that can reliably identify the most promising drugs to test could save time and resources in these screens, and expedite the treatment of these disorders.

Many have tried to build such a tool, using automated literature-parsing pipelines(8–10) and pathway-analysis algorithms that rely on authors’ interpretation of their own data, but there is increasing interest in approaches that instead utilize the data directly. RummaGene(11) parses data tables in the medical literature, as opposed to the main text, and links genes together based on their frequency of co-occurrence; All RNA-Seq and ChIP-Seq Sample and Signature Search (ARCHS4)(12) provides not only a database of uniformly computed read counts from the RNA-Sequencing (RNA-Seq) samples in GEO, but also correlations between two genes’ expression levels across human and mouse samples; and RummaGEO(13) automatically groups samples within each RNA-Seq study in ARCHS4’s read count database, calculates differential expression, and links genes together based on their frequency of co-occurrence in differentially expressed gene (DEG) lists. What these approaches have in common, alongside other correlative measures such as CoRegNet(14), gene set enrichment analysis(15), and the weighted gene co-expression network analysis(16), is that they do not establish temporality, and thus cannot distinguish causation from correlation.

In one attempt to address this deficit, RummaGEO was paired with known transcription-factor binding sites to infer this causation for transcription factors, in the ChEA-KG resource(17), formerly known as hGRN-ChEAR(18). Another attempt was the Connectivity Map (CMap)(19), which provides this information on temporality, in a massive screen of gene-perturbations and drug treatments in numerous human cell lines, using a microarray targeting 978 genes. CMap’s flagship study found molecules with similar effects on the transcriptome to the knockouts of specific tyrosine kinases, and experimentally verified that those molecules were binding to and inhibiting those kinases(19). It was unclear how many false predictions they had to sift through before they settled on these correct ones, and augmenting signatures over 10-fold from a small microarray might not be an adequate substitute for RNA-Seq(20). Still, the data have shown considerable promise in guiding drug-repurposing efforts, as Huang et al.(21) showed a strong correlation between similarity of drugs’ differential gene expression profiles (or transcriptomic “signatures”) and their propensity to have the same therapeutic indications. That same year, Novak et al.(22) used an AI-augmented prioritization system incorporating CMap’s microarrays, discrete pathway data, and manually curated RNA-seq data from MECP2-disruption studies to screen for therapeutics in a Xenopus laevis model of Rett syndrome (followed by a mouse study), and showed phenotypic improvements from the use of vorinostat. Similarly to CMap’s drug-screen, however, the drug was selected manually, not just by the AI. These approaches have considerable potential in optimizing treatment regimens and potentially identifying novel therapeutics, but for rare Mendelian diseases with no known therapy, it is unknown how (or whether) transcriptomic data can truly expedite the search for an effective treatment.

Thanks to data-sharing policies and centralized repositories such as the Gene Expression Omnibus (GEO)(23), RNA-Seq has become one of the most abundant omic data modalities. It has previously been proposed to match signatures of drug treatments and either disease states(24–26) or gene disruptions(24) to predict the therapeutic benefit of the given drug, but such approaches need to be scalable to the millions of samples in GEO. An expert on a specific gene could easily do a literature-search to find RNA-Seq studies disrupting that gene, or testing one specific treatment. However, if a gene has no already-known regulators, then in order to find potential candidates through signature-matching, the expert would have to repeat this painstaking literature search for thousands of genes and compounds. To make GEO-wide analyses more feasible, multiple labs ran uniform alignment on the raw reads of most of the human and mouse samples in the Sequence Read Archive (SRA, which is tied to the metadata in GEO) to convert them into gene (or transcript) read counts. This resulted in databases such as ARCHS4(12), Recount3(27), and Digital Expression Explorer 2 (DEE2)(28), which together lowered a tremendous barrier in the re-use of these data.

The remaining challenge, however, is identifying the experiments that were done and which samples were in which group. Descriptions of such are provided by GEO in plain language, but meta-analyzing numerous studies from different authors requires a means of encoding all of the different annotation formats into one. Artificial intelligence (AI) has previously been used to systematically encode the study descriptions in GEO, but none quite capture what we need in order to automatically collate gene-disruption and drug experiments for a meta-analysis. Meta-SRA(29) uses a rule-based text-reasoning graph to extract key study information, but is no longer maintained as of 2020. GEOracle(30) uses a support vector machine to separate samples, and GeMI(31) uses a transformer(32) model, but neither tool indicates what perturbagen distinguishes one group from the other, and the latter is no longer available. Lastly, the Crowd Extracted Expression of Differential Signatures (CREEDS) tool(33) uses crowd-sourced manual annotations of microarray studies in GEO (including what gene was being perturbed, what drug was administered, and which samples were in the experimental and control groups) to train a classifier to provide these annotations for other studies. After the classifier was run, the reusable training data and the predictions that it made on other studies were both made publicly available. However, it is not being updated; it was only trained and run on microarray studies; and it would appear that the participants in the crowd-sourced data curation annotated whichever studies they wanted to, which may bias the dataset in favor of the studies that are easy to interpret. The last point is not necessarily a drawback, as one could argue that the ambiguously annotated studies can add noise and should thus be avoided; but it is still worthwhile to attempt to salvage them.

Our aim is to fully operationalize the annotation of publicly available transcriptomic studies on drug-treatments and gene-disruptions, and determine whether these automatically curated studies can drive reliable regulatory relationship predictions that other tools would have missed. Hence, we introduce Signature-based Networks from Automatically Curated Knockout, Knockdown, and Small-molecule Studies (SNACKKSS), which uses a series of fine-tuned Bidirectional Encoder Representations from Transformers (BERT) models(34) to annotate the GEO metadata, and feeds these annotations directly into a signature-matching and relationship-prediction pipeline without any manual intervention. With this pipeline in place, we first evaluate its automated metadata curation, then test whether curated perturbation signatures can effectively prioritize correct modulators over incorrect ones, and finally ask whether SNACKKSS-based predictors add nonredundant value alongside existing predictive tools. Lastly, we demonstrate the vastly improved predictive ability achieved by an ensemble of many predictive tools (including a variant of SNACKKSS), which is available for download at snackkss.nrihub.org.

## Materials and Methods

### Reagents; Biological Resources: Not applicable Novel Programs, Software, Algorithms

The code used to run the analyses in this article is divided into four publicly available GitHub repositories: https://github.com/coledeisseroth/SNACKKSS_NLP, https://github.com/coledeisseroth/SNACKKSS, https://github.com/coledeisseroth/SNACKKSS_Eval, and https://github.com/coledeisseroth/SNACKKSS_Revision. The latest study curations and corresponding relationship predictions are available at https://snackkss.nrihub.org.

### Web Sites/Data Base Referencing

Gene Expression Omnibus(23), from no later than July 24, 2025. https://www.ncbi.nlm.nih.gov/geo/ CREEDS(33), accessed on December 16, 2023. https://maayanlab.cloud/CREEDS/

ARCHS4’s read counts(12), version 2.5. https://archs4.org/download

ARCHS4’s correlations(12), version 2.4. https://archs4.org/download Recount3’s read counts(27) on July 30, 2025. https://rna.recount.bio/

DEE2’s read counts(28), accessed from July 31 to August 1, 2025. https://dee2.io/. DEE2 underwent a major structural update after we accessed it, and the code we use to download and parse it does not work on the latest version. The version of DEE2 that we acquired is available upon reasonable request.

PARMESAN(10), corpus timestamped on February 15, 2025. https://parmesan.nrihub.org/#/home PubTator3(9), accessed July 20, 2025. https://www.ncbi.nlm.nih.gov/research/pubtator3/

Connectivity Map(19), level-5 MODZ scores, downloaded on July 20, 2025, but some of the data files are time-stamped at December 16-17, 2020. https://clue.io/data/CMap2020#LINCS2020

DGIdb(35), version from December 6, 2024. https://dgidb.org/downloads

Reactome’s functional interaction dataset(36), version from April 14, 2025. https://reactome.org/download-data

DIOPT(37), accessed July 25, 2025. https://www.flyrnai.org/cgi-bin/DRSC_orthologs.pl Entrez gene database, accessed July 20, 2025. https://www.ncbi.nlm.nih.gov/gene/ PubChem Substance IDs, accessed July 19, 2025. https://pubchem.ncbi.nlm.nih.gov/

Guide to Pharmacology(38), timestamped at June 15, 2026. https://www.guidetopharmacology.org/ PharmGKB (merged with ClinPGX)(39), accessed June 1, 2026. https://www.clinpgx.org/ ChEMBL(40), accessed June 1, 2026. https://www.ebi.ac.uk/chembl/

Drug Repurposing Hub(41), accessed April 29, 2026. https://repo-hub.broadinstitute.org/repurposing#home TRRUST v2(42), timestamped on April 15, 2018. https://www.grnpedia.org/trrust/

### Random selection of high-throughput sequencing series

To ensure an unbiased sampling of GEO studies, we queried GEO on May 20, 2024 for all series IDs under the label “Expression profiling by high-throughput sequencing”, of which there were 97, 822. We shuffled them with a random seed of “2024”. C.A.D., a predoctoral trainee, annotated the first 10 of these series, indicating any relevant genetic or pharmacologic perturbations and how to distinguish control from experimental samples. Under his supervision, a high-school-level researcher, B.B., annotated another 615 during a three-week internship. C.A.D. subsequently made corrections in order to ensure that all of the annotated studies were accounted for (either fully processed or recognized as not processable) by the scripts that were processing them—these scripts would be reading “soft” files instead of the GEO web portal that the manual curators were using. He also changed any labels that he disagreed with. For transparency, we provide both the uncorrected and the corrected versions of this manually curated dataset (Supplemental tables 1 and 2, respectively), and for the remainder of this study, we only use the corrected version. We call our manually curated dataset “SNACKKSS-Manual Curation”, or “SNACKKSS-MC”, and display in Supplemental figure 1 the number of GEO series with each type of perturbation in it and in CREEDS’ dataset. Types of correction made by C.A.D. are plotted in Supplemental figure 2. In order to assess the stability of our labels, another high-school-level researcher, B.S.S., annotated the same 615 studies that B.B. did, indicating for which ones C.A.D. was consulted for guidance. These annotations are provided in Supplemental table 3, and we refer to them as “SNACKKSS-Second Curation”, or “SNACKKSS-2C”. Details on the incorporation of CREEDS’ dataset are provided in the Supplemental methods, and a statistical comparison between SNACKKSS-MC and SNACKKSS-2C is provided in Supplemental table 4.

**Table 1:**
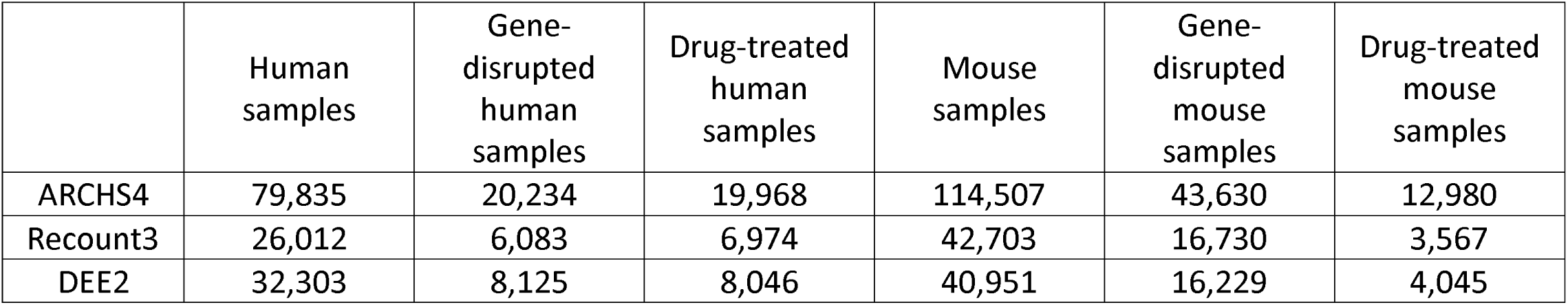
Perturbed samples identified by SNACKKSS.

**Table 2:**
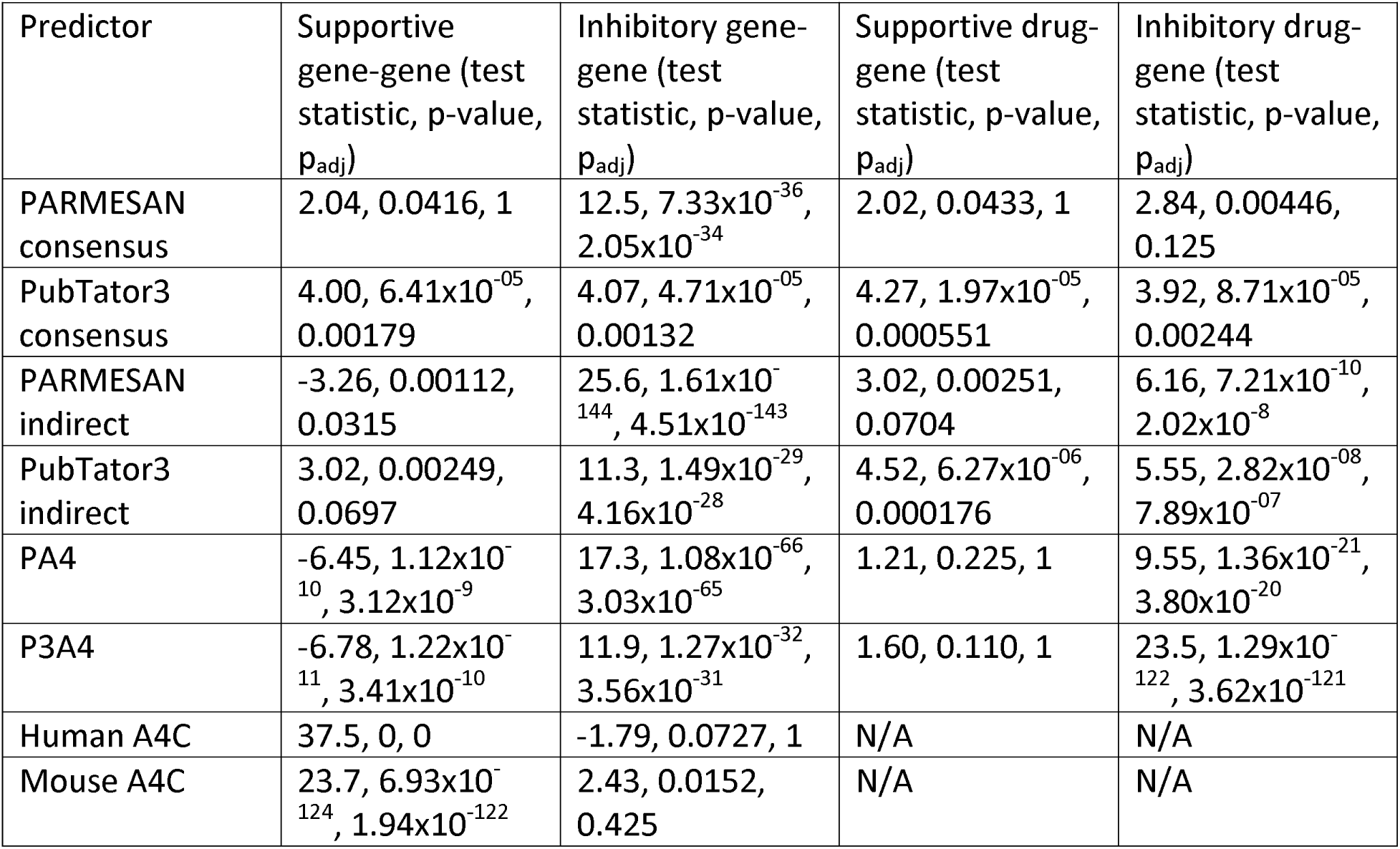
Performance of non-signature-based relationship predictors.

| Predictor | Supportive gene-gene (test statistic, p-value, $p_{adj}$) | Inhibitory gene-gene (test statistic, p-value, $p_{adj}$) | Supportive drug-gene (test statistic, p-value, $p_{adj}$) | Inhibitory drug-gene (test statistic, p-value, $p_{adj}$) |
| --- | --- | --- | --- | --- |
| PARMESAN consensus | 2.04, 0.0416, 1 | 12.5, $7.33 \times 10^{-36}$, $2.05 \times 10^{-34}$ | 2.02, 0.0433, 1 | 2.84, 0.00446, 0.125 |
| PubTator3 consensus | 4.00, $6.41 \times 10^{-05}$, 0.00179 | 4.07, $4.71 \times 10^{-05}$, 0.00132 | 4.27, $1.97 \times 10^{-05}$, 0.000551 | 3.92, $8.71 \times 10^{-05}$, 0.00244 |
| PARMESAN indirect | -3.26, 0.00112, 0.0315 | 25.6, $1.61 \times 10^{-144}$, $4.51 \times 10^{-143}$ | 3.02, 0.00251, 0.0704 | 6.16, $7.21 \times 10^{-10}$, $2.02 \times 10^{-8}$ |
| PubTator3 indirect | 3.02, 0.00249, 0.0697 | 11.3, $1.49 \times 10^{-29}$, $4.16 \times 10^{-28}$ | 4.52, $6.27 \times 10^{-06}$, 0.000176 | 5.55, $2.82 \times 10^{-08}$, $7.89 \times 10^{-07}$ |
| PA4 | -6.45, $1.12 \times 10^{-10}$, $3.12 \times 10^{-9}$ | 17.3, $1.08 \times 10^{-66}$, $3.03 \times 10^{-65}$ | 1.21, 0.225, 1 | 9.55, $1.36 \times 10^{-21}$, $3.80 \times 10^{-20}$ |
| P3A4 | -6.78, $1.22 \times 10^{-11}$, $3.41 \times 10^{-10}$ | 11.9, $1.27 \times 10^{-32}$, $3.56 \times 10^{-31}$ | 1.60, 0.110, 1 | 23.5, $1.29 \times 10^{-122}$, $3.62 \times 10^{-121}$ |
| Human A4C | 37.5, 0, 0 | -1.79, 0.0727, 1 | N/A | N/A |
| Mouse A4C | 23.7, $6.93 \times 10^{-124}$, $1.94 \times 10^{-122}$ | 2.43, 0.0152, 0.425 | N/A | N/A |

### BERT models were trained on manual annotations

Our processing pipeline had four key steps: (1) Study classification: whether the study is testing a gene disruption; (2) Sample classification: whether a given sample is receiving a gene disruption; (3) Target classification: what gene was disrupted; (4) Control classification: given one sample that had a specific gene disruption and one that did not, whether the latter is an appropriate control to the former. For example, if there are multiple cell lines tested, then the controls for a given disruption should come from the same line.

Knockout and knockdown experiments (KO and KD) are both accepted as gene-disruptions, but not overexpression (OE, which would have the opposite effect to disrupting it) or other modulations (OM, which will have unpredictable effects). We call all four experiments “gene perturbations”, but we only refer to KO and KD as “gene disruptions”. Drug-treatment studies are curated analogously to this, and we also refer to them as a type of perturbation, but not a gene perturbation.

We tested three BERT(34) models on each of these tasks: DistilBERT(43), a smaller BERT model trained to mimic the original one; BioBERT(44), the original BERT model, fine-tuned on medical literature; and BioMedBERT(45), the original BERT model trained exclusively on the medical literature, from default weights instead of BERT’s pre-trained ones. The Supplemental methods elaborate on how we introduce these classification tasks to the BERT models. For each model, we would measure the precision, recall, and F1 score in each task. We used 4-fold cross-validation (4FCV, see **Supplemental table 5** for the series IDs put into each group, and the Supplemental methods for further details on how we conducted it) to train and test each model on both SNACKKSS-MC and CREEDS’ manually curated dataset. We also tested the performance from training each model on CREEDS’ dataset followed by SNACKKSS-MC, as well as SNACKKSS-MC followed by CREEDS’ dataset. Because SNACKKSS-MC is a random selection of high-throughput sequencing studies, instead of human-selected microarrays, we used the model that performed the best on SNACKKSS-MC, even if another model performed better on CREEDS’ dataset.

We calculated standard (which we call “unsmoothed”) precision (true positives / (true positives + false positives)), recall (true positives / (true positives + false negatives)), and F1 scores (2 * precision * recall / (precision + recall)), as well as smoothed versions of these metrics that are more conservative. Smoothed precision is calculated as true positives / (true positives + false positives + 1). Smoothed recall is true positives / (true positives + false negatives + 1). The smoothed F1 score is calculated as 3 * smoothed precision * smoothed recall / (smoothed precision + smoothed recall + 1). The denominators are incremented by 1 to penalize smaller sample sizes, and the F1 score formula is adjusted so that it remains scaled from 0 to 1. Smoothed precision and recall can be as low as zero (with zero true positives), and can get infinitely close to 1 as the number of true positives approaches infinity. Therefore, their ranges are both [0, 1). With smoothed precision and recall values of zero, the smoothed F1 = 3*0*0/(0+0+1) = 0, and as smoothed precision and recall approach 1, the smoothed F1 score approaches 3*1*1/(1+1+1)=1. Therefore, the range of the smoothed F1 score is also [0, 1). Further details on input-formatting, accuracy testing, and synonym resolution are provided in the Supplemental methods.

### The best fine-tuned model is identified for each classification task

For each task—classifying studies, perturbed samples, targets, and then controls—we test each BERT model after training it on either our dataset (SNACKKSS-MC), CREEDS’ dataset, SNACKKSS-MC and then CREEDS’, or CREEDS’ and then SNACKKSS-MC. 4FCV allows us to test each fine-tuned model on the full dataset. For each classification task (study, sample, target, and control) and each perturbation type (gene-disruption or drug), we identify the apparatus (which starting BERT model to use, which training corpora to use, and in what order) that had the highest smoothed F1 score on classifying SNACKKSS-MC’s studies through 4FCV. We then train that starting model on the entire training corpus (as opposed to three quarters of it as we do in 4FCV), which yields the finalized model for that task and perturbation type (there are eight models in total). For gene-disruptions and drugs, we run the study classifier on all of the human and mouse RNA-Seq study descriptions that we were able to acquire; then the sample classifier on the sample descriptions from all studies labeled as positive by the study classifier; then the target classifier on the descriptions of all samples labeled as positive. We then run the control classifier on the samples with automatically identified targets. Control classification is more computationally intensive than the other steps, because we are classifying pairs of samples, of which there can be millions in a single study. We thus apply five filters to the sample pairs (see Supplemental methods) to make the job feasible for our machines.

When training BioMedBERT on the first epoch of SNACKKSS-MC (after having trained it on CREEDS’ dataset), we assess whether the training data saturated this model. We measure this saturation along two dimensions: growth (whether the performance improves throughout training) and diminishing returns (whether the improvement progressively declines), and we define “saturation” as either the lack of steady growth or the presence of diminishing returns. Detailed mathematical definitions are provided in the Supplemental methods.

### Perturbation signatures are calculated across curated experiments

Because different studies can use different RNA-Seq alignment tools to establish read counts for each gene, the standard approach to RNA-Seq meta-analyses is to re-align the raw reads, using a uniform alignment pipeline(12, 27, 28, 46). This can be a trivial task for two or three studies, but when conducting large-scale meta-analyses across hundreds of thousands of samples, the computational cost of alignment requires dedicated resources. Multiple research teams have taken on this role, generating publicly available resources such as ARCHS4(12), Recount3(27), and DEE2(28), making it cost-effective and reproducible to calculate signatures for the tens of thousands of gene-disruption and drug-studies. In each of the three pre-computed read count datasets, ARCHS4, Recount3, and DEE2 (specifically, DEE2’s Kallisto(47) expression files), we procure any available read counts for that sample and its controls. Our approach to handling samples from different read count datasets is described in the Supplemental methods.

Having normalized each sample to its respective controls, we next need to establish overall differential gene expression patterns across samples that had the same gene disrupted. There are many established ways to collapse signatures into unified consensuses—CMap uses a “MODZ” score consisting of a weighted average of expression z-scores in perturbed samples(19), and Marino et al.(20) tested multiple consensus approaches for their ability to detect CMap drug signatures that are consistent with manually curated signatures from GEO. Future work may include determining the optimal collapsing method for finding regulatory relationships. For this study, however, we simply calculate the z-score of the perturbed samples’ z-scores (which we call the “nested z-score”), which should suffice if signature-matching is a valid predictive measure. For a disrupted gene A, we gather the samples that had A disrupted, and for each such sample X we take X’s control population and calculate the mean and standard deviation of every measured gene B’s read count, in transcripts per million (TPM). We then calculate a Z-score for X’s expression of B, by subtracting the control mean from X’s TPM read count for B, and dividing the difference by the control standard deviation. This gives an expression Z-score for gene B in each gene-A-disrupted sample. We calculate the mean and standard deviation of the z-scores of gene B expression across all of the samples that had gene A disrupted. Lastly, we calculate a z-score from this distribution (nested z-score), thus giving one unified estimate of the effect of gene A disruption on gene B expression. Drug signatures are calculated analogously, where A is an administered drug instead of a disrupted gene.

### Perturbation signatures are matched to predict regulatory relationships

The remaining question is whether the correlation between the signatures of two perturbations is suggestive of their relationship. Theoretically, two genes with similar disruption signatures would play roles supporting each other(19), whereas two genes whose disruptions have opposite effects on the transcriptome would have functions that normally oppose each other. Likewise, a small molecule whose administration yields similar or opposite effects to a gene’s disruption would oppose or support that gene’s function, respectively. To match signatures, we use a formula of our own design, which we give the moniker “Differential F1” or “DF1”. This formula is commutative, and can be run at high throughput in Bash. There are numerous existing methods for signature-matching, such as the weighted connectivity score(19), nemoCAD(22), or simply a Pearson or

Spearman correlation (both of which have been used in the context of co-expression(12, 48), and the latter of which has been used for matching drug signatures(21)). These correlation metrics are not feasible to run at the scale that SNACKKSS reaches, but since Huang et al.(21) established the Spearman correlation as a strong precedent for signature-matching, we provide a streamlined benchmark between it and DF1. The Supplemental methods describe how we compare them to each other, and explain why using a Spearman correlation would lead to an impractically long runtime. Future work will include finding the optimal signature-matching method, but we use DF1 for a high-speed proof of concept due to the number of signature pairs that need to be matched in this study. If signature matching is a viable relationship-prediction method, DF1 should be sufficient, even if it is not optimal.

DF1 is calculated as follows, for matching the signatures of Perturbations A and B: we first set a threshold T, where we will only accept a differentially expressed gene (DEG) if the absolute value of its nested z-score is larger than T. We then determine the ability of A’s signature to “detect” the DEGs in B’s signature, and quantify this with a smoothed-F1-like score. Let STP (supportive true positives) = the number of DEGs shared by A and B in the same direction (i.e. both z-scores are the same sign), OTP (opposing true positives) = the number of shared DEGs where A and B yielded opposite directions, FP (false positives) = the number of DEGs in A’s signature but not B’s, and FN (false negatives) = the number of DEGs in B’s signature but not A’s. We then calculate supportive and opposing smoothed precision (SP and OP, respectively) as:

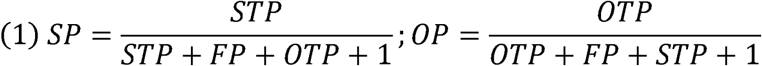

Likewise, we calculate supportive and opposing smoothed recall (SR and OR, respectively) as:

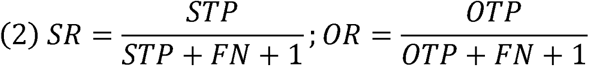

And we calculate the supportive and opposing smoothed F1 scores (SF1 and OF1, respectively) as:

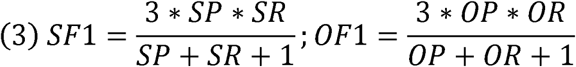

Our final directionality score, like the one used by PARMESAN(10), penalizes the presence of conflicting information in both directions:

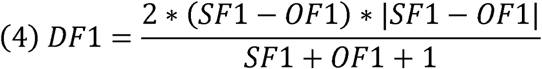

The 1s added to the denominators (which we call “smoothing”) are designed to penalize instances where there are few genes in a signature.

We then set the threshold T through leave-one-out cross-validation (LOOCV): we calculate all DF1 matches with T = 0, 0.1, 0.2… up to 0.9. For each relation in the manually curated dataset from modifier A to target B, we determine which of the ten thresholds is the most effective at prioritizing correct predictions over incorrect ones (highest log-rank test statistic) among the relations involving neither A nor any of its targets. Using that threshold, we then determine the DF1 score between A and B. We run separate LOOCV optimizations for positive and negative prediction scores, and eliminate any predictions where the directionality is different between the two optima. In other words, for a given two entities, if the positive LOOCV optimization led to a positive prediction score, and the negative optimization led to a negative one (or vice versa), the relationship between those two entities will not be predicted.

### SNACKKSS variations assess whether the chosen features are optimal

One could argue that any failure in SNACKKSS’ predictive ability is due to poorly conducted signature-matching, rather than inherently poor performance of signature-matching. Therefore, we test its performance after three different variations, each of which challenges an assumption we make in building this pipeline. Testing these variations alone still leaves open the possibility that a combination of them, and possibly additional ones, would lead to stronger classification performance. For computational feasibility, however, we evaluate the effect of each variation alone relative to our baseline.

The first variation, which we call “Target-down”, only accepts gene-disruption samples if the supposed target gene had decreased expression compared to the control samples (drug treatment data are kept as they are). This accounts for the possibility that the tool’s performance is hampered by studies where either the methodology, the labeling, or the curation was improper. The second variation, “ARCHS4-only”, only uses samples from ARCHS4 (the largest of the three read count datasets), rather than incorporating Recount3 and DEE2. Ideally, all read counts should be calculated using the same version of the same alignment tool, so limiting to one challenges our assumption that it is safe to conflate signatures calculated on different platforms. The third variation, “Mouse”, uses mouse data instead of human. KO/KD data are more than twice as abundant in mouse tissue as in human (see Results), so the former may offer more robust predictions.

### Indirect predictions of regulatory relationships are derived from signature-matches

A major contribution offered by PARMESAN is its ability to predict undiscovered regulatory relationships by linking the known ones together. We have demonstrated its ability to expand our repertoire beyond what was explicitly written in the literature(10), and the same principle may apply to SNACKKSS and bolster its predictive ability.

Notably, gene coexpression networks are a common means of identifying pairs of genes that may have similar or related functionality(14, 16). They do not establish temporality—when two genes have increased expression, we do not know whether one increase caused the other. However, they offer a far larger repertoire of potential relations than the literature- or signature-matching-based approaches do. ARCHS4 provides a readily accessible table of Pearson correlations between the expression of any two genes (covering 29, 082 human and 24, 530 mouse genes), across all of the human and mouse samples that they curated. The correlations are a powerful predictor of supportive gene-gene relations (see Results), so we investigated whether they could improve the literature- and RNA-based relationship predictors. We first linked PARMESAN’s gene-gene relation consensuses to either its own consensuses (as PARMESAN normally does to make indirect predictions), or to said correlations. Our linking formula, “PA4”, simply treats ARCHS4’s correlation coefficients as gene-gene relationship consensus scores, which we feed into the I_CA_ formula as described in Deisseroth et al.(10).

Similarly, we test the performance of linking SNACKKSS’ DF1 scores to ARCHS4’s correlations, treating the former the same way we treat PARMESAN’s consensus scores, and call the resulting predictor “SA4”. We evaluate the performance using the default setup for SNACKKSS, as well as with the three aforementioned variations, Target-down, ARCHS4-only, and Mouse.

### The Connectivity Map is fed into the same relationship prediction pipeline

As we did with SNACKKSS, we tested the ability of the Connectivity Map (CMap) to correctly identify regulatory relationships. Specific details on our processing of CMap data are provided in the Supplemental methods. In addition to testing a default prediction pipeline, we introduce four variations that challenge unique assumptions we make regarding our approach. The first, “Inferred”, uses all of the quantified gene expression levels, instead of just the landmarks. The second, “shRNA only”, excludes CRISPR KO data when calculating gene-disruption signatures.

The third we call “MCF7”, which only uses samples in the MCF7 cell line, and tests the effect of taking the tissue type into consideration. Although SNACKKSS is implicitly trained to select control samples that are of the same tissue type as a perturbed sample, it does not consider tissue type when calculating consensus signatures—which, one could argue, might hinder its performance. Quantifying the comparability of samples from different studies would require not only a strict encoding of plain-text tissue descriptions in GEO, but also a ground truth for defining two samples as comparable to each other. CMap, however, does have a strict encoding of the tissue used. Hence, we tested the performance from using only samples from their most-tested cell line, MCF7.

The last variation we call “OE-corrected”. Correlations between the signatures of two perturbations can reflect their relationship, but they can also be due to a cellular stress response or off-target effects. Overexpression studies would theoretically be a promising solution to this conundrum: DEGs that change in the same direction in response to a KO/KD or OE of Gene A are likely unrelated to the altered level of A, but if the KD and OE change a gene’s level in opposite directions, that DEG is far more likely to truly be responding to A’s level, thus giving us a cleaner and more faithful signature(49). Unfortunately, per both our curation and that of CREEDS, OE studies are less common than KO and KD, but with CMap’s repertoire of OE data for 3, 542 target genes (2, 753 of which also have either KO or KD data), we may be able to test this hypothesis. Hence, this final variation only uses target genes that have both OE and KO/KD data, takes the average magnitude between the two, and excludes any DEGs that changed in the same direction.

### Other predictive tools were evaluated

Ultimately, we have the following candidate predictive tools:

“SNACKKSS”: Using the signature-matches from SNACKKSS (default setup) to predict regulatory relationships. Positive DF1 scores are interpreted as supportive gene-gene and inhibitory drug-gene relations, and negative scores are interpreted as inhibitory gene-gene and supportive drug-gene relations.

“CMap”: Using the signature-matches from the Connectivity Map (default setup) to predict regulatory relationships, as we do with SNACKKSS.

“PARMESAN consensus”: Using the consensus scores from PARMESAN, as we have done previously(10). “PARMESAN indirect”: Using the indirect relationship predictions from PARMESAN, as we have done previously(10). Also referred to as PARMESAN’s indirect predictions.

“PubTator3 consensus” Feeding PubTator3’s extracted regulatory relationships into PARMESAN’s consensus formula.

“PubTator3 indirect”: Feeding PubTator3’s extracted regulatory relationships into PARMESAN’s consensus formula, and feeding those consensuses into the indirect prediction formula. Also referred to as PubTator3’s indirect predictions.

“Human A4C”: Using the Pearson correlations from ARCHS4’s human gene coexpression matrix to predict gene-gene relations. Positive and negative correlation coefficients are interpreted as predictions of supportive and inhibitory gene-gene relations, respectively. We also refer to this dataset as just “A4C”.

“Mouse A4C”: Same as “Human A4C”, except using ARCHS4’s mouse gene coexpression matrix instead. “SA4”: Linking the signature-matches from SNACKKSS to A4C, as described above.

“CMA4”: Linking the signature-matches from CMap to A4C, analogously to SA4.

“PA4”: Linking PARMESAN’s consensuses to A4C, instead of to its own gene-gene relationship consensuses. “P3A4”: Linking the consensuses from PubTator3 to A4C, similarly to PA4.

### Redundancy of SA4’s predictions is assessed

SA4 is outperformed by another tool for every prediction task (see Results). Its poorer precision and recall, however, do not necessarily mean that it is not useful in an ensemble of predictive tools. It draws relationships from a data source nigh untouched by PARMESAN and PubTator3, and can identify relations that would have been missed when using the literature alone. Nevertheless, incorporating a noisy predictive tool that sometimes looks accurate through pure chance can ultimately hinder the prioritization of candidate drugs (see **Table 3** for examples). Therefore, our final assessment of this tool’s utility asks whether SA4 is helpful or harmful when added to an ensemble of other predictors.

**Table 3:** Examples of beneficial and detrimental effects of incorporating SA4.

| Modifier | Target | Direction, per DGIdb | Source | Prediction without SA4 | Confidence without SA4 (0-1) | Prediction with SA4 | Confidence with SA4 (0-1) |
| --- | --- | --- | --- | --- | --- | --- | --- |
| Indisulam | GAPDH | inhibitor | Guide To Pharmacology | N/A | N/A | Inhibitory | 0.989796 |
| Dinoprostone | RHOC | activator | Guide To Pharmacology | N/A | N/A | Inhibitory | 0.991453 |
| Panobinostat | SIRT2 | inhibitor | Targeted Agents in Lung Cancer | Supportive | 0.343348 | Inhibitory | 0.98913 |
| ORY-1001 | GEM | activator | Guide To Pharmacology | Supportive | 0.224781 | Inhibitory | 0.956318 |

Our setup is akin to LOOCV: For a given drug A known to modulate a given gene product B, we exclude any relations involving A or B from DGIdb, resulting in the database DGIdb_-AB_. For each predictor (PubTator3, P3A4, PARMESAN, PA4, CMA4, and SA4, as well as Human and Mouse A4C for gene-gene relations), we measure the smoothed precision of the scores above any given threshold in predicting the directionality of the relations in DGIdb_-AB_, thereby mapping its directionality scores to smoothed precision estimates *p̂* (which we also call “confidence”). Then, when predicting the relation between A and B, each of the predictors will provide a putative directionality and a *p̂*. The directionality we posit is that of whichever predictor gave the highest *p̂*, and we score the prediction with that *p̂*. The resulting precision, P, is the percent of left-out A-to-B relations that were correctly predicted when we only accepted predictions with *p̂* above a given threshold.

This simulation represents a user’s approach to using multiple tools to predict a relation that has not been discovered: if none of the tools make a confident prediction, the candidate drug will have low priority, and if SA4 is redundant with the other predictors, then it will not confidently predict any “undiscovered” relationships that the other tools could not predict more confidently (where confidence is defined as ^). This simulation also accounts for the imperfection of our reference databases: SA4 might yield overly confident predictions that appeared strong amongst the relations covered by DGIdb, but ultimately assign the wrong directionality to relations that PubTator3 and P3A4 were not confident about, thus worsening the overall performance. Again, we compare the recall with versus without SA4 using a binomial test for the proportion of precision levels at which we were able to achieve higher recall by adding SA4 (with an expected rate of 0.5)— but we emphasize that this test statistic has few practical implications, as the best model to use depends on the precision one is willing to accept.

### Statistical analyses

Statistical methods are described in their appropriate sections. All p-values are two-sided. When comparing the performance of our BERT classifiers to one another, we did not use any statistical tests, because our goal was to decide which model to use, not to determine whether the differences in performance could occur by random chance. One could potentially test for significantly different performance using two-proportion Z-tests for precision and recall, but our goal is to decide which model to use, and significance does not affect this model-selection decision. While DistilBERT is faster than the other two models, the tasks in this pipeline are feasible for all three, so we would use BioBERT or BioMedBERT if it performed even slightly better than DistilBERT.

When testing whether the BERT models outperformed random guessing, we assumed that a baseline model would randomly guess positive or negative for any entity it classifies, with a 50% chance of each. For this predictor, we calculated the expected precision; and used a binomial test to compare it to the observed precision of the BERT model. We ran the same statistical test to compare the recall to that of the same random predictor, but we note that this test is not as informative, because a random predictor can easily achieve a favorable recall value at high statistical significance, simply by having a low threshold. The same cannot be said for precision. Further details on this analysis are provided in the Supplemental methods.

We display both smoothed and unsmoothed precision, recall, and F1 scores, as described above. This smoothing is similar to Bayes-Laplace Smoothing (BLS), except that BLS brings all proportions closer to 50%, which penalizes small sample sizes for proportions above 50% while rewarding them if they are below 50%. This more-stringent version of BLS, by not incrementing the number of successes, penalizes small sample sizes regardless of the proportion. 95% confidence intervals around percentages are calculated using a binomial distribution via Microsoft Excel’s BINOM.INV function. For smoothed and unsmoothed F1 scores, 95% confidence intervals are calculated by measuring the binomial 95% confidence interval (same formula) around the smoothed and unsmoothed (respectively) precision and recall.

All p-values given are uncorrected unless stated otherwise. We used Bonferroni correction (i.e. multiply all p-values by the number of hypotheses tested) whenever there was a collection of hypotheses, where we would consider any one success to be an overall success. A rationale of each multiple-hypothesis correction is provided in the Supplemental methods.

When measuring the proportion of SNACKKSS’ alleged gene-disruption samples for which the supposed target gene decreased in expression (**Supplemental table 9**), we correct the binomial p-value for 6 hypotheses: 3 precomputed read count datasets, each covering 2 species. Similarly, the binomial p-values measuring whether the CMap-based gene-perturbation signatures predominantly show increased or decreased expression of the target gene are corrected for 9 hypotheses: 3 types of perturbation: KO, KD, and OE; multiplied by 3 categories of measured gene: landmark, best-inferred, and inferred.

To determine whether the signature-based relationship predictors (using SNACKKSS or CMap) prioritize correct predictions over incorrect ones **(Supplemental table 10),** we correct the log-rank p-values for 72 hypotheses: 2 (direct signature-matching, and linking those signature-matches to the coexpression matrix from ARCHS4) by 2 (supportive and inhibitory relations) by 2 (gene-gene and drug-gene relations) by 9 (4 variations of SNACKKSSS and 5 of the CMap-based pipeline). We similarly correct the log-rank tests for the prioritization ability of non-signature-based tools (Table 2) for 28 hypotheses: 8 gene-gene + 6 drug-gene relationship predictors = 14 predictors; each predicting 2 directions, up- and down-regulators.

For our ablation test, these binomial p-values are corrected for 36 hypotheses: 10 gene-gene and 8 drug-gene relationship predictors ablated; multiplied by 2 directions, supportive and inhibitory. Lastly, when comparing DF1 to a Spearman correlation **(Supplemental table 19),** Bonferroni correction covers 3 matching formulae (DF1, a Spearman correlation, and a Spearman correlation requiring a significant p-value) by 2 directions (support and inhibition) by 2 prediction types (gene-gene and drug-gene relations), resulting in 24 hypotheses.

## Results

### Manual GEO curators predominantly agree with each other

We manually annotated GEO series to train and test our automated study-curation pipeline, generating the “SNACKKSS-MC” dataset; and an additional curator re-annotated the same series for a point of comparison, generating “SNACKKSS-2C”. Among the studies in SNACKKSS-2C that were annotated independently, the labels predominantly agreed with those from SNACKKSS-MC. The number of labels agreed upon for each classification task is provided in Supplemental table 4, alongside rates of agreement, relevant statistics, and 95% confidence intervals. Overall, the two curated datasets agreed on 92.2% of gene-disruption and 91.7% of drug-study labels (binomial p values undetectably low). Given that they agreed on a study label, they agreed on 95.7% of samples receiving that disruption and 94.2% of samples receiving a drug (binomial p values undetectably low). Given that they agreed on a gene-disrupted or drug-treated sample, they had unsmoothed F1 scores of 0.960 and 0.860 in detecting each other’s target labels, respectively (p-values could not be calculated). For a given agreed-upon gene-disrupted or drug-treated sample, they showed 71.4% and 93.2% weighted agreement on control samples, respectively (binomial p-values undetectably low). Without this weighting, they agreed on 52.4% (p undetectably low, worse than the expected agreement) and 96.0% (p undetectably low), respectively, warranting an error analysis. We provide in the SNACKKSS_Revision GitHub repository a list of studies where the gene-disruption-control disagreement occurred, and an explanation of why. Typically, one curator found a field to stratify the samples by, that the other curator did not.

Overall, these agreement statistics support the use of SNACKKSS-MC as a training and benchmarking set for large-scale prioritization, but they should not be interpreted as evidence that every individual sample-level annotation is correct. Accordingly, the automatically curated labels from SNACKKSS should be treated as a scalable resource for hypothesis generation rather than as a ground truth.

### BERT-based metadata classification shows high accuracy despite hardware-driven noise

We evaluated the performance of each BERT model trained on each manually curated corpus (SNACKKSS-MC or CREEDS’ dataset, or both), for each of the eight classification tasks: study, sample, target, and control classification for gene-disruption and drug-treatment experiments. Supplemental Table 6 displays the accuracy metrics from running the pipeline on a 40- and a 64-core CPU. We also track the progressive improvement of BioMedBERT on a 64-core CPU as it is trained on CREEDS’ dataset followed by SNACKKSS-MC, in Supplemental table 7, with the 4-fold cross-validation performance on SNACKKSS-MC plotted in Supplemental figure 3. When measuring model saturation, we observed that SNACKKSS-MC saturated all of the models except for the drug-study and drug-target classifiers. This resulted from using a training batch size of 1; with a batch size of 16, gene-disruption study, drug study, drug target, and gene-disruption control classification were not saturated, and the other four tasks were. It is unsurprising that the larger batch size led to less-saturated models, given that fewer steps of training are done in that epoch.

Importantly, the two CPUs did not yield the same results when running the same models on the same data, and they did not always agree on which model performs the best. To test whether this variability was due to differences in hardware architecture, or a software-level discrepancy that we had neglected, we ran a small portion of the NLP pipeline (see Supplemental methods) on another 64-core CPU. The prediction scores it produced were identical to those yielded by the first 64-core CPU, and not to those from the 40-core device. This indicates that the difference in output is a direct result of differing hardware architecture. It has previously been shown that devices with different numbers of cores can generate different output in the same machine-learning tasks(50), and the authors attributed the phenomenon to the differing levels of parallelism. While they were not specifically testing CPUs, the principle holds regardless: imperfect floating-point precision coupled with multiplications not being done in the exact same order will inevitably lead to rounding errors accumulating differently, thus resulting in different output.

With different machines performing differently, we needed to assess the impact on classification performance. Figure 1A shows examples of stable (gene-disruption study) and unstable (drug-treatment sample) classification performance. For the former, the models’ unsmoothed F1 scores were always less than 2% different between CPUs, while for the latter, they could be as much as 18% different. However, the unsmoothed F1 scores of the respective top-performing models from each CPU were always less than 2% apart. This led us to ask whether, if we let each CPU select and run its top-performing model through leave-one-out cross-validation (LOOCV), they ultimately converge on similar performance. Figure 1B shows the resulting unsmoothed F1 scores achieved by the two CPUs, and the differences between them could range from 0.00406% to 17.5%. The supporting data, with and without smoothing, are in Supplemental table 8. This instability is important to be aware of, but it does not preclude the use of BERT models for automated curation, because the reasoning of deep-learning models is difficult to trace regardless, and since the classifications are saved, any downstream prediction errors can be traced back to the BERT curation if need be. Furthermore, regardless of the CPU used, the unsmoothed precision achieved by the best model in each task (through the aforementioned LOOCV) was significantly better than what one could achieve through random guessing (Supplemental table 8, with all binomial p-values undetectably low). This means that the NLP models run on either machine will preferentially detect true information in these studies, more so than could be explained by hardware-level noise. Hence, there is still merit in testing the ability of the saved output to guide relationship predictions.

**Figure 1:**
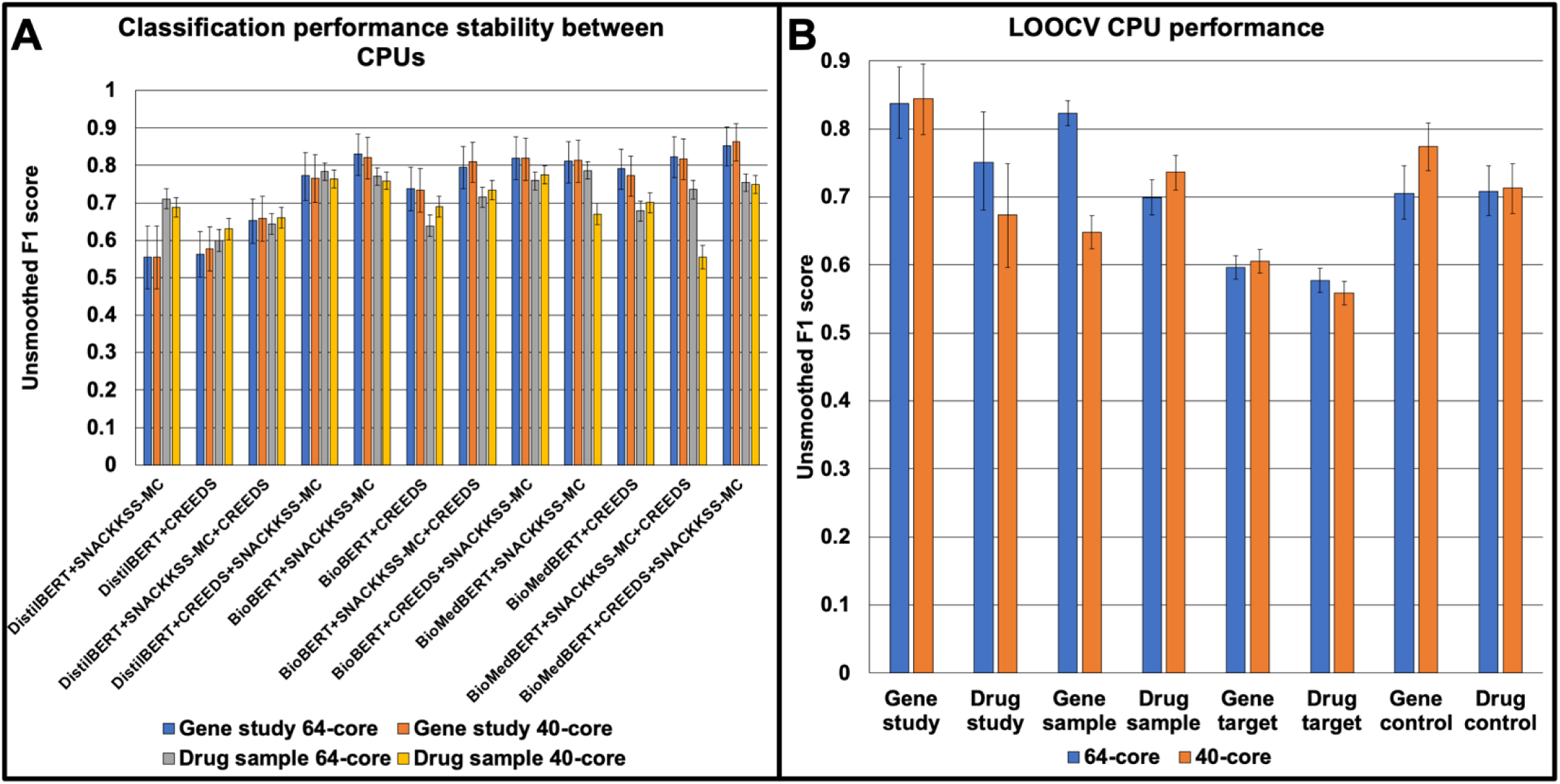
BERT models can perform differently on different machines. (A) Through 4-fold cross-validation, we trained DistilBERT, BioBERT, and BioMedBERT on either SNACKKSS-MC or CREEDS’ dataset, or both (in both orders) on a 64-core and a 40-core CPU, for gene-disruption study and drug-treatment sample classification, and tested their performance on SNACKKSS-MC. We measure the performance as an unsmoothed F1 score. (B) We have each machine identify and use its optimal model (highest smoothed F1 score) for each task through leave-one-out cross-validation (LOOCV), and the resulting unsmoothed F1 scores can differ between them, from less than 1% to more than 17%. The error bars are calculated by measuring the binomial 95% confidence interval around a predictor’s unsmoothed precision and recall values, and calculating the unsmoothed F1 scores resulting from the minimum and maximum values.

Using the 40-core CPU, our final gene-disruption study curation pipeline used BioMedBERT trained on SNACKKSS-MC alone for gene sample and drug target classification, BioBERT trained on CREEDS’ dataset followed by SNACKKSS-MC for drug samples, and BioMedBERT trained on CREEDS’ followed by SNACKKSS-MC for the other five tasks. To give one successful example, this pipeline accurately annotated GSE160415 (per the 4-fold cross-validation training BioMedBERT on our dataset alone on the 40-core CPU). This study tested a knockout of Mark3 in mouse osteoblasts. The gene- and drug-study classifiers marked this study as positive and negative, respectively. The sample classifier identified GSM4873066, GSM4873067, and GSM4873068 (and no other samples) as receiving a gene KO/KD; and the target classifier exclusively procured “Mark3” instances from their descriptions. For each KO sample, the control classifier marked all of the wild-type samples as controls—although this particular study had no candidate controls that were not in fact controls. Meanwhile, this pipeline mislabeled the study GSE224420 (testing ROR2 knockdown in a cancer cell line(51)) as non-gene-disruption. These examples are meant to illustrate the type of information extracted by SNACKKSS. Indeed, we cannot trace the reasoning of these labels, but we can trace downstream errors back to the labels themselves.

For each pre-computed read count dataset, we provide the total number of samples available through that dataset that were identified by SNACKKSS to be part of a gene-disruption or drug experiment, as well as the number of samples that were specifically identified as gene-disrupted or drug-treated. We count human and mouse samples separately.

### Gene expression changes are largely concordant with SNACKKSS’ study annotations

Having established pipelines for curating gene-disruption and drug studies, we next measured the value of interpreting these automatically curated studies. We ran our two pipelines on all of the qualified studies, and identified which samples had read counts in ARCHS4, Recount3, and DEE2 (**Table 1**). We found that across all read count datasets, the supposedly disrupted genes predominantly decreased in expression in both humans and mice. Numbers of samples with increased and decreased target gene expression are plotted in **Supplemental figure 4** and listed in **Supplemental table 9**, alongside the binomial 95% confidence interval around each percentage. The corrected binomial p-values were all undetectably low for human ARCHS4 (81.0%), mouse ARCHS4 (73.6%), human Recount3 (83.3%), mouse Recount3 (72.5%), human DEE2 (78.7%), and mouse DEE2 (71.9%). Granted, the expression-increases ranged considerably higher than the decreases, likely because expression levels can go to infinity, but not below zero. Ultimately, disrupted genes were far more likely to show decreased expression than increased, which suggests that they and their controls were effectively identified.

### Matching disruption signatures yields weak predictions of gene modulators

We tested the high-speed signature-matching algorithm, Differential F1 (DF1), for its ability to predict the directionality (support or inhibition) of the manually curated regulatory relationships provided by Reactome (functional interactions where one protein supports or opposes another(36)) and DGIdb (The Drug-Gene Interaction Database, indicating molecules that support or oppose specific gene products(35)). We call these datasets “manually curated databases”, or “MCDBs”, and use them as a gold-standard. Smoothed precision across score thresholds is plotted in Figure 2A, with log-rank test statistics for all signature-matching-based predictors provided in **Supplemental table 10** and raw correct/incorrect counts provided in **Supplemental table 11. Figure 2** uses smoothed precision, while **Supplemental figure 5** plots the same data using unsmoothed precision instead. To give one example of a successful prediction, DGIdb asserts that gefitinib, an epidermal growth factor receptor inhibitor(52) (Also known as ZD1839 or “Iressa”, PubChem SID 532631), opposes Human Epidermal Growth Factor Receptor 2 (HER2, also known as ERBB2, Entrez ID 2064), and there are multiple indirect mechanisms through which it is theorized to do so(53). Using the relationship predictions involving neither gefitinib nor any of its targets, 0.3 was the best DEG z-score threshold for prioritizing correct inhibitory relations over incorrect ones, and 0.9 was the best for prioritizing supportive relations. Using 0.3 as the threshold, across the gefitinib treatment and ERBB2 disruption studies (in human tissue), there were 9, 991 DEGs that went in the same direction (STP), 9, 797 in the gefitinib but not the ERBB2 disruption signature (FP), 20, 159 in the ERBB2 disruption but not the gefitinib signature (FN), and 4, 283 shared DEGs that went in opposite directions (OTP). With the above formulae, DF1 = 0.0428, suggesting that gefitinib yields a similar effect on the transcriptome to disruption of ERBB2, and likely opposes it. Using 0.9 as the threshold, STP = 405, OTP = 222, FP = 2, 625, FN = 21, 424, and DF1 = 3.36×10^−5^—a weaker match, but in agreement with the inhibition-optimized match, hence we keep our prediction. This is only one example of matching signatures coinciding with the relationship between those two entities, and we must evaluate whether the principle holds true across the regulatory relationships in Reactome and DGIdb.

**Figure 2:**
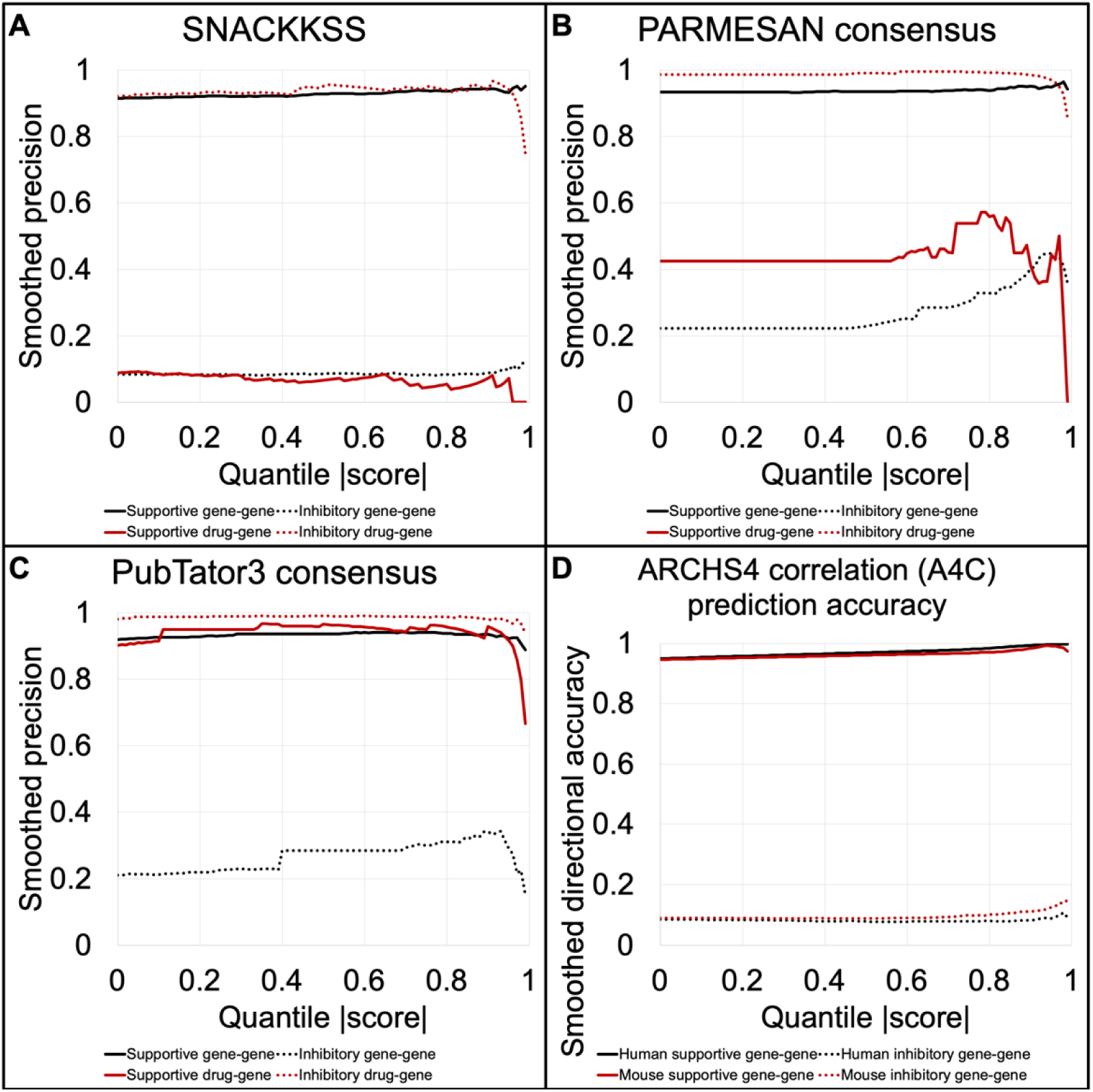
Whole-signature matching is a weak predictor of regulatory relationships. We measure the smoothed precision (likelihood of getting the correct directionality, up-versus down-regulation, with the denominator incremented by 1) among the relationships posited by SNACKKSS (A), PARMESAN’s consensuses (B), PubTator3’s consensuses (C), or ARCHS4’s expression correlations (D). The X axis is the minimum score quantile above which we accept putative relations—for example, at 0.9, we accept only the top-10%-scoring candidates. The manually curated databases (MCDBs) we use as a gold standard are Reactome (gene-gene) and DGIdb (drug-gene relations). If a prediction tool is effectively prioritizing correct relations over incorrect ones, its smoothed precision should increase with higher scores (and, by extension, score quantiles).

Before taking directionality into consideration, the gene-gene relations seemed to be prioritized well (all log-rank test statistics > 6), while the drug predictions were not (all < 1.7, **Supplemental table 10**). However, we observed a shared bias between SNACKKSS and the MCDBs. Positive matches between perturbations (which suggest supportive gene-gene and inhibitory drug-gene relations) are more common than negative ones, with stronger matches being more likely to be positive (limiting to the signature-matches involving known modulators in Reactome and DGIdb, log-rank p < 10^−22^ for both gene-gene and drug-gene matches, at all DEG thresholds except 0.9 for drug-gene relations, where p = 0.141). Reactome contains 100, 724/106, 656 (94.4%, 95% CI 94.3-94.6%) supportive relations, and DGIdb contains 18, 616/25, 313 (73.5%, 95% CI 73.0-74.1%) inhibitory relations. To address this bias, we analyzed positive match scores separately from negative ones. If shared directional bias is the sole contributor to accuracy improvements, then when we guarantee a fixed directionality of our predictions, the supportive and inhibitory relations catalogued in the MCDB should decline at the same rate as the score threshold increases. Therefore, this separation can assess whether an increase in the magnitude of the score truly improves accuracy.

We provide the log-rank test results (in the format of “test statistic, p-value, p_adj_”) for the literature-based tools and ARCHS4’s gene expression correlations (“A4C”) in their ability to prioritize correct regulatory relations from Reactome and DGIdb over incorrect ones. “p_adj_” is the Bonferroni-corrected p-value, with 28 hypotheses. “Indirect” refers to linking gene-gene or drug-gene relationship consensuses from a given database to gene-gene relation consensuses from that same database, using PARMESAN’s indirect prediction formula I_CA_. “PA4” and “P3A4” refer to the linking of PARMESAN’s and PubTator3’s consensuses (respectively) to the gene expression correlations in ARCHS4, also using the I_CA_ formula. “N/A” means not applicable— expression correlations alone cannot predict drug-gene relations.

Unsurprisingly, the prioritization was notably weaker after this separation (log-rank p = 1.49×10^−4^ for supportive gene-gene, and 1 for inhibitory gene-gene, supportive drug-gene, and inhibitory drug-gene relations). For PARMESAN, this was not as severe of an issue (Figure 2B, log-rank test results provided in Table 2). However, this finding with SNACKKSS informed us that, even for PARMESAN, we must consider the two directions separately when we estimate precision, as scores of -1 and +1 are not necessarily equally reliable.

Existing resources can provide analogous regulatory relationship predictions that can be tested in the same way. PubTator3’s extracted gene-gene and drug-gene relations can also be fed into PARMESAN’s consensus formula, and they demonstrate strong accuracy (Figure 2C). Furthermore, ARCHS4’s gene coexpression matrix can be used for the same purpose, where a stronger correlation coefficient is interpreted as a higher-priority prediction. From comparing ARCHS4’s gene expression correlations in human tissue to the known regulatory relationships in Reactome, strong positive correlations were a decisive indicator of supportive gene-gene relations (Figure 2D). The negative correlations, however, were not as robust: there was no significant prioritization of inhibitory relations from human data, and although there was in mice, the unsmoothed precision in identifying them never reached 15%, which we do not expect most users to find acceptable—thus, one might gravitate toward PARMESAN for this purpose instead. Intuitively, each prediction task is performed best by a different algorithm.

### Compared to human data, mouse data yield better identification of gene-gene relations

To assess for flaws in our approach to signature-based relationship predictions, we tested the performance of SNACKKSS after implementing three methodological variations: “Target-down”, which only accepts gene-disruption samples if the supposedly disrupted gene had decreased expression compared to the controls; “ARCHS4-only”, which only uses pre-computed read counts from ARCHS4; and “Mouse”, which uses mouse data instead of human. We plot the unsmoothed precision and recall of each variation of SNACKKSS (Target-down, ARCHS4-only, and Mouse) in Supplemental figure 6, with the log-rank test statistics for prioritization of correct over incorrect in Supplemental table 10 and correct/incorrect counts in Supplemental table 11. None of the variations led to strong prioritization of drug-gene relations (the best result was from the Target-down approach, p_adj_ = 0.226 for inhibitory and 1 for supportive relations), nor inhibitory gene-gene relations (with the best performance from Target-down, p_adj_=1).

### ARCHS4’s expression correlations can bolster other predictive tools

Due to the strong performance observed from using ARCHS4’s co-expression matrix to predict supportive gene-gene relations, we asked whether linking them to literature-based consensuses offered a better precision-recall balance. The correct/incorrect counts across score quantiles for PARMESAN-and PubTator3-derived indirect predictions, and for those derived from linking PARMESAN or PubTator3 to ARCHS4’s expression correlations (PA4 and P3A4, respectively), are in **Supplemental table 12**, with log-rank statistics provided in **Table 2**. When plotting precision against recall (both unsmoothed, **Figure 3**), P3A4 offered record-breaking recall across precision levels for finding inhibitory drugs.

**Figure 3:**
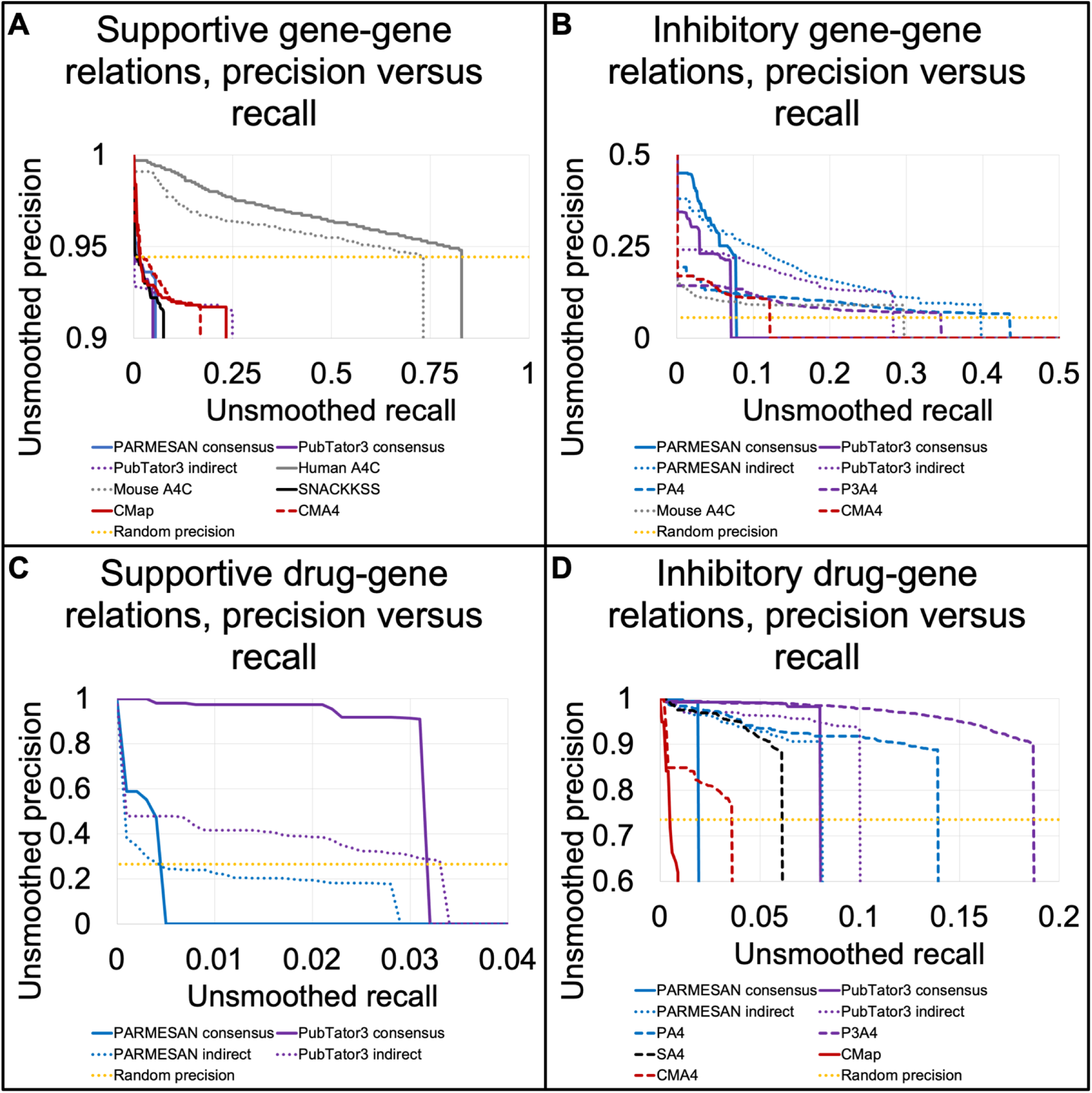
Signature matches synergize with coexpression data and contribute to negative drug-gene relationship identification. We display the performance of different predictive tools in identifying supportive gene-gene (A), inhibitory gene-gene (B), supportive drug-gene (C), and inhibitory drug-gene (D) relations. Specifically, we plot unsmoothed precision (Y axis) against unsmoothed recall (X axis) in identifying manually-curated-database (MCDB) relations, and mark the precision that one would achieve from random guessing (equal to the percent of the MCDB that matches the predicted direction). The predictors are PARMESAN’s consensuses, indirect predictions, and ARCHS4-linked predictions (“PA4”), PubTator3’s consensuses, indirect predictions, and ARCHS4-linked predictions (“P3A4”), gene expression correlations from ARCHS4’s human and mouse datasets (“Human A4C” and “Mouse A4C”, respectively), signature-matches from SNACKKSS or Connectivity Map (“CMap”), and ARCHS4’s correlations linked to signature matches from SNACKKSS (“SA4”) or Connectivity Map (“CMA4”). We only plot a predictor if it prioritizes correct predictions over incorrect ones, with an uncorrected log-rank p < 0.05.

When we linked SNACKKSS’ gene-gene DF1 scores to ARCHS4’s correlations (“SA4”, with log-rank prioritization statistics in Supplemental table 10, raw correct/incorrect counts across score thresholds in Supplemental table 13, and unsmoothed precision and recall plotted in Figure 3), we only observed strong prioritization of inhibitory drugs (log-rank p = 8.85×10^−28^). Among the drug-gene relations verified by DGIdb, the highest-scoring inhibitory relationship from SA4 was from indisulam (PubChem Substance ID 12015040) to GAPDH (Entrez ID 2597), with a z-threshold set as 0.2 through LOOCV, resulting in a score of 3.58. DGIdb agreed that this relationship was inhibitory, citing the Guide to Pharmacology(38). The single strongest contributor to this prediction (with the other contributors provided in **Supplemental table 14**) involved similar signatures of indisulam treatment and disruption of PPM1G (Entrez ID 5496, DF1 = 0.183), a gene whose expression has a correlation coefficient of 0.503 with that of GAPDH. Note that this one correct prediction is only meant to serve as an example, not as evidence for or against the performance of this method. The global comparison to DGIdb, however, does suggest effective prioritization of inhibitory substances.

As we did with direct signature-matching (“SNACKKSS”), we measured the effect of limiting SA4 to samples with down-regulated targets, limiting to ARCHS4, and using mouse data (full performance statistics are in **Supplemental table 10**, correct/incorrect prediction counts across score thresholds are in **Supplemental table 13**, and unsmoothed precision and recall are plotted in **Supplemental figure 6**). Mouse data improved the gene-gene relationship predictions (p_adj_ = 0.00168 and 0.0638 for supportive and inhibitory relations, respectively), nothing salvaged the supportive drug-gene relations, and nothing sharply improved the inhibitory drug-gene relations. Future work may include using cross-validation to select and evaluate the optimal method, which would require additional streamlining due to the runtime-multiplicity of testing different combinations of features. For this study, we adhere to the default regardless of the outcomes observed, and compare the variations purely for investigational purposes. Beyond the use of mouse data, this test did not identify any obvious ways to improve SNACKKSS’ performance.

### Connectivity Map data can also fuel signature-based relationship predictions

SNACKKSS uses public RNA-Seq data to predict regulatory relationships, but it is not the first systematic, large-scale application of the signature reversion paradigm. The Connectivity Map(19) (CMap) is a massive series of microarrays measuring the levels of 978 genes in multiple human cell lines perturbed with thousands of compounds, shRNAs, CRISPR knockouts, and overexpression vectors. Despite the limited breadth of the microarray, the database provides all of the features that we have been trying to automatically annotate, in a machine-readable format, as well as the cell line, which SNACKKSS does not definitively account for. Because their database can be run through the same prediction pipeline that SNACKKSS uses, we cannot say that SNACKKSS makes a novel contribution until we know whether the same contribution could be made when matching signatures from CMap instead.

CMap imputes the expression of over 11, 000 genes, and its definition of “best inferred” is that the predicted expression level has a statistically significant correlation with the actual level. However, a statistically significant correlation is not the same as a reliable one. To assess the usability of these inferences, we first measured the differential expression of the targeted genes in the shRNA, CRISPR KO, and overexpression samples, in comparison to the control samples (control vector, control vector, and DMSO, respectively) from the same cell line. For the landmark genes, there was decreased expression of 84.7% (p_adj_ undetectably low) of the CRISPR targets, 84.6% (p_adj_ undetectably low) of the shRNA targets, and 33.4% (p = 9.98×10^−206^) of the overexpression targets (**Supplemental figure 7A**). For the best inferred genes, these percentages were 54.2% (p_adj_ undetectably low), 48.2% (p = 4.00×10^−25^), and 46.2% (p = 1.95×10^−21^), respectively (**Supplemental figure 7B**), and for the inferred genes, they were 51.0% (p_adj_ = 0.192), 46.7% (p = 8.34×10^−07^), and 44.4% (p = 3.58×10^−05^), respectively (**Supplemental figure 7C**). The sample counts supplying Supplemental figure 7, as well as the binomial 95% confidence intervals around the percentages listed above and uncorrected binomial p-values, are provided in **Supplemental table 9**. The strong concordance between the perturbation of landmark genes and their change in expression suggests that the direct measurements were accurate and the gene-perturbations were usually effective. Given this, the relative lack of concordance seen from imputed gene expression levels suggests, as expected, that they are not nearly as reliable as the direct measurements.

We next determined whether matching signatures from CMap led to accurate predictions of regulatory relationships. Using DF1, we compared KO/KD signature matches to Reactome relations, and compound-to-KO/KD matches to DGIdb relations (log-rank prioritization statistics are in Supplemental table 10, correct/incorrect counts are in Supplemental table 11, and unsmoothed precision and recall are plotted in **Figure 3**). Upon matching signatures, we observed decent prioritization of supportive gene-gene relations (log-rank p = 6.56×10^−4^ for supportive gene-gene, 1 for inhibitory gene-gene, 0.0184 favoring the wrong direction for supportive drug-gene, and 0.0701 for inhibitory drug-gene relations).

As we did with the other predictors, we determined whether the DF1-based predictions from CMap could be bolstered when linked to the expression correlations from ARCHS4 (“CMA4”, log-rank prioritization statistics are in Supplemental table 10, correct/incorrect counts are in Supplemental table 13, and unsmoothed precision and recall are plotted in Figure 3). This linking led to improved prioritization of inhibitory gene-gene relations (Log-rank p = 1.55×10^−6^ for supportive and 1.59×10^−4^ for inhibitory relations).

Among the default CMap-derived predictor and the variations, four and three effectively prioritized supportive and inhibitory gene-gene relations, respectively; and one and seven effectively prioritized supportive and inhibitory drug-gene relations, respectively. Correct and incorrect counts across score thresholds are provided in Supplemental table 13, log-rank statistics are in Supplemental table 10, and unsmoothed precision and recall are plotted in Supplemental figure 8). This demonstrated some merit to limiting to a specific tissue type, and—to our surprise—using inferred expression levels.

Adding a new predictive tool to one’s repertoire can add valuable new information alongside unwanted noise. We provide examples of both resulting from using the SNACKKSS/ARCHS4 hybrid (“SA4”) alongside the other predictive tools. Each predictor scores its prediction of a relationship’s directionality, and we use the Drug-Gene Interaction Database (DGIdb) relations not involving the modifier nor the target in question to calculate the expected smoothed precision ( ^, or “confidence”) of the predicted relation, given the score that it received. We believe whichever model has the highest ^, and show what the prediction (and the highest ^) would have been had we included or excluded SA4. Indeed, SA4 can add both correct and incorrect relationship predictions to our repertoire, and can reverse the predicted directionality of formerly correct and incorrect predictions, meaning that we must directly measure whether the benefits outweigh the drawbacks— in other words, whether including SA4 truly improves our recall without sacrificing precision.

### SA4’s inhibitory drug predictions are non-redundant with other tools

SA4 alone is not the strongest predictor in any regard. Since it uses a unique source of information, however, it might find regulators that the other tools never would have considered, and could thus be a beneficial addition to one’s repertoire. Nonetheless, including a new predictor in a decision process can be helpful or harmful (**Table 3**). To test what SA4 adds beyond what other, more-powerful predictive tools do, we use LOOCV to estimate the smoothed precision ( ^) of each predictor, and use the most confident prediction (highest ^) for the left-out relation.

We determined the precision and recall achieved when using all predictors, and when using all of them except one. To provide a simplified account of the effects of adding each predictor, **Table 4** indicates whether doing so increased the maximum precision and recall. This is by no means a comprehensive description of the benefits of each predictor, as benefiting one facet would often come at a cost to the other. Users should ultimately decide which set of models to use based on their own preferred balance of precision against recall, and we plot the two (unsmoothed) upon removing each predictor in **Figure 4** (drug-gene relations) and Supplemental figure 9 (gene-gene relations), derived from **Supplemental table 15**. The fraction of precision thresholds for which each predictor improved recall is in **Supplemental table 16**, with 95% confidence intervals around the percentage, binomial test statistics, and corrected and uncorrected p-values. Individual predictors’ precision/recall distributions in the LOOCV test can be derived from **Supplemental table 17**. We did not test the performance of removing multiple predictors, as our goal with this test is to showcase the tradeoff (or the strict benefit or burden) of each predictor. And indeed, each one usually did introduce such a tradeoff—but we again see A4C’s strong contribution to the identification of supportive gene-gene relations, and P3A4’s benefit in inhibitory drug prediction.

**Figure 4:**
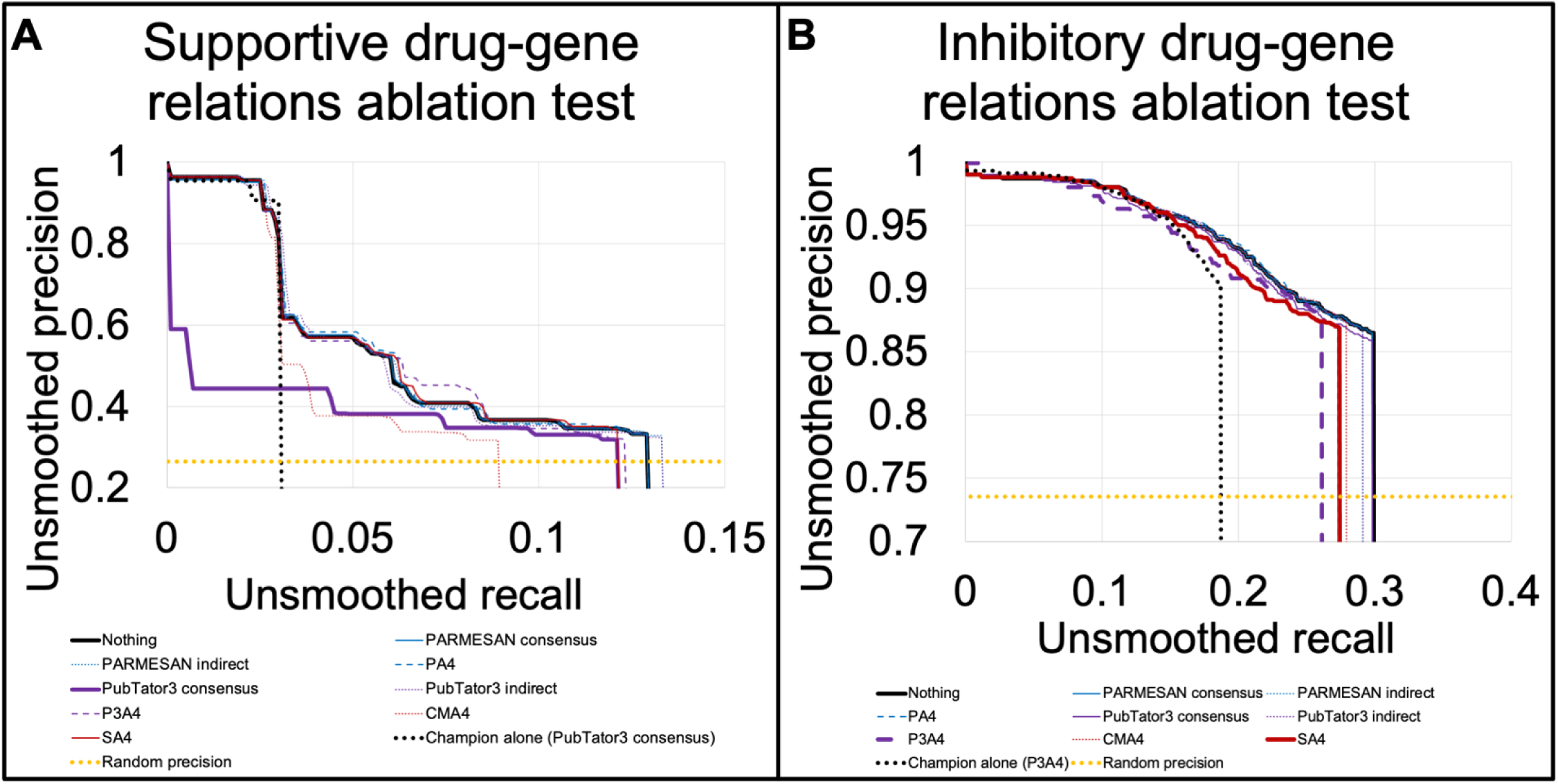
Contribution of each predictor to drug prediction. Through leave-one-out cross-validation (LOOCV), we evaluated the use of all predictors in the ensemble to identify previously-unseen drug-gene relations. Here, we test the contribution of each predictor to such an effort by removing it from the repertoire and measuring how the performance changes. We plot unsmoothed precision and recall in identifying supportive (A) and inhibitory (B) drug-gene relations, and mark the “Random precision”, or the precision that one would achieve from random guessing (equal to the percent of DGIdb relations that match the predicted direction). Whether a predictor is helpful usually depends on the precision one is willing to accept. Hence, depending on where a user sets the cutoff, some predictors—even if they do well on their own—will do more harm than good when used alongside more powerful tools. SA4 yields improved recall across most precision levels in finding inhibitory drug-gene relations, and for drug prediction overall, the ensemble of predictors yielded a better precision-recall balance than the reigning champion alone.

**Table 4:** Overview of the benefits of each predictor.

|  | Supportive gene-gene | Inhibitory gene-gene | Supportive drug-gene | Inhibitory drug-gene |
| --- | --- | --- | --- | --- |
| **Human A4C** | Both | Neither | Recall | Recall |
| **Mouse A4C** | Recall | Both | Recall | Recall |
| **CMA4** | Recall | Neither | Recall | Recall |
| **SA4** | Recall | Neither | Recall | Recall |
| **PARMESAN consensus** | Recall | Neither | Precision | Recall |
| **PARMESAN indirect** | Recall | Recall | Neither | Recall |
| **PA4** | Recall | Neither | Neither | Recall |
| **PubTator3 consensus** | Recall | Neither | Both | Recall |
| **PubTator3 indirect** | Recall | Neither | Neither | Recall |
| **P3A4** | Recall | Neither | Recall | Recall |

Adding SA4 was highly beneficial for predicting inhibitory drug-gene relations (outperforming recall at 394/413 precision levels, p_adj_ undetectably low), but not supportive ones (outperforming recall at 10/112 precision levels), nor gene-gene relations (outperforming recall at 207/2563 precision levels for supportive relations, and 1/15 for inhibitory). Furthermore, the ensemble of these predictors led to vastly improved recall in identifying drug-gene relations, compared to using the best predictors alone (PubTator3’s consensuses for supportive relations, and P3A4 for inhibitory ones), while for supportive gene-gene relations, it was comparable to using just ARCHS4’s correlations.

Unexpectedly, for inhibitory gene-gene and supportive drug-gene relations, removing datasets from this LOOCV evaluation sometimes led to more predictions. Indeed, our system is designed to believe whichever model is the most confident in its selected direction, and with all models inherently being more confident in one direction than the other, they can serve as a filter for candidate relations in the rarer direction. In other words, all models more confidently predict supportive than inhibitory gene-gene relations, so in order for this LOOCV system to predict an inhibitory relation, none of the models must confidently say it is supportive. For inhibitory gene-gene relations, this seemed to detract from the effort, as using PARMESAN’s indirect predictions alone yielded overall stronger performance than the ensemble. This phenomenon could be an indication to train a sophisticated machine-learning classifier on the scores assigned by different models (as others have done for their ensemble predictors(21)), and future work may include exploring some of the infinitely many possible implementations.

Since we know that SA4 makes a unique contribution to our overall predictive effort, we sought to answer the question, for a given gene product that one is studying, how much have we improved the likelihood that one will be able to find a promising inhibitor of it? We plot the number of genes for which we were able to correctly identify an inhibitor (which we call the target “coverage”) across accepted precision levels, before and after adding SA4 to our repertoire (Supplemental table 18 and Supplemental figure 10). For example, if one desires 95% unsmoothed precision, SA4 allows one to target 40 more genes than one could without it, underscoring its benefit to drug-repurposing efforts.

### Performance evaluation against a Spearman correlation

DF1 was chosen as the signature-matching metric because of its high speed compared to more intensive correlation metrics, such as the Spearman correlation. However, since the latter is a precedented approach to signature-matching(21), a comparison of runtime and predictive accuracy is needed. We tested the performance of SNACKKSS and SA4, using a Spearman correlation instead of DF1 to match perturbation signatures. The Supplemental methods explain why a Spearman correlation is impractical in this setting, and how we streamlined our test to accommodate it. Supplemental table 19 shows the log-rank statistics for each predictor prioritizing correct relations over incorrect ones, as well as the number of correct relations identified at each smoothed and unsmoothed level of precision. Meanwhile, Supplemental figure 11 plots the unsmoothed precision and recall achieved by each approach. While the Spearman correlation yielded lackluster results (log-rank p_adj_ > 0.48), requiring its p-value to be below 0.05 led to robust inhibitory drug predictions from SA4 (p_adj_=0.000108), to a similar degree that we could achieve using DF1 (p_adj_ = 0.00669). This would suggest that severely downweighing (as done by DF1) or outright excluding (as done when we filter the Spearman correlations) the weak correlations allows for cleaner and more accurate predictions made by SA4.

### Performance evaluation on an unused gold standard

Since we have used Reactome and DGIdb before to test the performance of PARMESAN, we needed to test the performance of these tools on an independent gold standard. Reactome and DGIdb have not been updated with new relations since we finalized this version of SNACKKSS, but there are other datasets we can use. For gene-gene relations, there is a dataset called TRRUST v2(42), a manually curated corpus of 5, 050 transcription-factor/target relations (4, 195 excluding Reactome’s relations), which specifies whether the target is being up- or down-regulated. For drug-gene relations, the Drug Repurposing Hub(41) provides 11, 529 relations, excluding those from DGIdb. There are also the publicly available datasets that DGIdb used to pull from, which have been updated since DGIdb’s last release. This includes PharmGKB(39), Guide to Pharmacology(38), and ChEMBL(40); which have 179, 3, 314, and 589 relations (respectively) that are not present in DGIdb. Altogether, these four datasets provide 15, 516 drug-gene relations not registered in DGIdb. For each predictor, we ran log-rank tests to measure whether it was prioritizing correct relations over incorrect ones (**Supplemental table 20**), and plotted precision against recall **(Supplemental figure 12, Supplemental table 20)**. We also re-ran the ablation test with this new gold standard **(Supplemental figure 13, Supplemental table 21)**.

As we had previously observed, the ensemble predictor struggled with gene-gene relations. Its prioritization was still effective (log-rank p = 0.000145 for supportive and 1.11×10^−7^ for inhibitory relations), but it typically did not improve recall at better-than-random precision. As we had also previously seen, though, it vastly improved our ability to find regulatory drugs at better-than-random precision, and effectively prioritized correct predictions over incorrect ones (log-rank p = 3.61×10^−8^ for supportive and 1.69×10^−37^ for inhibitory drugs).

Meanwhile, SA4’s contribution to the ensemble’s overall precision-recall curve for finding inhibitory drugs was not as decisively beneficial with the new gold-standard dataset, but it did improve recall at better-than-random precision. Furthermore, SA4 continues to prioritize correct pharmacologic inhibitors over incorrect ones (log-rank p = 1.83×10^−5^), meaning that its benefit primarily lies in supplementing our repertoire with modulators that other tools did not know about, and are still far more promising than a random guess. Testing these tools against a new gold-standard dataset supports the benefit of the ensemble predictor in identifying drug-gene relations, and clarifies that of SA4 in finding inhibitory drugs.

For each predictor on each task (supportive and inhibitory gene-gene and drug-gene relationship prediction), we display whether incorporating its predictions (through LOOCV) led to a higher maximum unsmoothed recall, a higher maximum unsmoothed precision, both, or neither. ARCHS4’s correlations (Human and Mouse A4C) could not be used for drug prediction, hence the corresponding fields are labeled as “N/A”.

## Discussion

There are tens of thousands of rare genetic disorders that still lack targeted therapies, but are not studied widely enough that we can identify promising candidate regulators. The millions of publicly available RNA-Seq samples offer tremendous promise in guiding drug discovery for these conditions, and many researchers have developed innovative ways to make them findable, accessible, interoperable, and reusable(54); but there has been one missing step required for feeding them into fully automated drug prediction: a machine-readable registry of gene-disruption and drug-treatment studies. SNACKKSS fulfills this role and carries it directly into the effort to identify molecules that can up- or down-regulate specific gene products. While it is not the best solitary predictor in any regard, we demonstrate that the predictive approach SA4 gives high priority to correct inhibitory drugs, and that adding it to our repertoire substantially improves our recall in finding them.

We frame SNACKKSS (and the other drug-prediction tools) in the context of targeting Mendelian disease genes, but pharmacologically targeting specific proteins underlies much of modern medicine—inhibiting oncogenes in cancer, or clotting factors in coagulopathies; or substituting a deficient hormone in an endocrine disorder. SA4 correctly identified Gleevec (Imatinib) as an inhibitor of ABL1 at 97% confidence, and this is a part of the standard of care for chronic myeloid leukemia with a BCR-ABL fusion(55). It also correctly identified estradiol as an inhibitor of RANKL at over 92% confidence(56), consistent with its known implications in treating osteoporosis(57). We have not proven that any of these tools can guide therapeutic decisions, and their output should never be interpreted as medical advice, but even the more common, multifactorial disorders can often benefit from modulation of single proteins. Moreover, a Mendelian haploinsufficiency disorder that has no way to pharmacologically activate the deficient protein could be targeted indirectly by pharmacologically inhibiting a gene product that opposes it. MicroRNA-132 (miR-132) is known to inhibit MECP2 translation(58), and SNACKKSS’ raw drug signatures indicate that lithium and ibuprofen decrease the level of this microRNA (Nested Z-scores = -5.79957 and -2.67735, respectively). We are not aware of any evidence that these agents have a therapeutic role in treating Rett syndrome, but if one is conducting a drug screen for MeCP2-supporting compounds, they might be valuable candidates to include. As is illustrated by this example, however, selecting modulators that inhibit an inhibitor requires a clear understanding of what other gene product needs to be modulated. It nonetheless highlights the importance of automated GEO study curation: an expert could easily compile a registry of studies testing one specific drug or gene-disruption, and would likely annotate them more accurately than SNACKKSS; but when searching for modulators that have never been explicitly tied to a gene product of interest, the process of meta-analyzing other perturbations (such as drugs that we did not know modulate the level of miR-132) must be automated.

Along with our findings, we offer four key resources to the public, which are available at snackkss.nrihub.org. The first is a database of automatically curated RNA-Seq experiments from GEO testing gene disruptions and drug treatments, detailing the control samples that correspond to each perturbed one, as well as what perturbation was administered. As with any automatically curated dataset, the accuracy is not perfect, and these labels should not be taken as a ground truth. Nonetheless, they will likely be an asset to users who want to systematically gather previous experiments testing any of a broad list of drugs or gene disruptions.

The second is the consensus signatures of each gene disruption and drug treatment (searchable by both perturbagen and differentially expressed gene), and the third is a database of predicted regulatory relationships, automatically constructed from the SNACKKSS and SA4 approaches. However, we provide the disclaimer that we only expect the inhibitory drug predictions from SA4 to offer a valuable contribution to the identification of novel protein regulators. The fourth is the ensemble of the different predictors we tested (for download only, and only for drug-gene relations), which displays each tool’s prediction and confidence, and which we expect to be a particularly strong advance in drug prioritization.

While SNACKKSS offers a systematic implementation of the signature reversion paradigm, it is not a definitive evaluation of the principle itself. There are major limitations in this study—first and foremost, the signature that a treatment is reversing has to be part of the disease’s pathogenicity(48), rather than part of a compensatory effect. Quantifying this distinction will be difficult, but doing so would massively improve the utility of this type of predictor, especially for translational purposes.

Additionally, our redundancy tests are not comprehensive, and even the inhibitory drug predictions might be overshadowed by some configuration of other predictive tools. We have also previously mentioned that our precision metric for relationship prediction does not account for the possibility that there is no relationship between two entities, and instead assumes that there is either support or inhibition(10). Furthermore, although we have tested variations of the signature-matching pipeline, we cannot claim any aspect of our pipeline to be optimal. If we simply adopt variations as we find that they improve performance, we run the risk of overfitting our methodology to the gold-standard upon which we test it—so future work will include exploring alternatives systematically but prudently.

There are also two minor nuances to mention: first, we have operated at the sample level, normalizing each experimental sample to its controls(19). One could instead operate at the experiment level, by grouping control and treatment samples with RummaGEO’s K-means clustering (13), calculating differential expression with DESeq2(59), and collapsing experiments with a random-effects meta-analysis(60, 61). We call this a minor nuance because, in contrast to the variations that we tested, this is simply another way to collapse the same data into one hypothesis, which should only make a substantial difference if one of the two methods is severely dysfunctional. These slower but more-intricate approaches could be worth testing, with some efficiency-optimization and all of the same due caution against overfitting.

Second, our metrics for precision and recall of the BERT classifiers could be overestimated, as we were not blinded to the manually curated data while we were building the input mechanism. The variability in performance from running the models on different machines also limits the strength of any claim about the exact NLP performance metrics. Nonetheless, the downstream analyses use saved classification output, allowing errors to be traced back to the curation stage when necessary.

For best practice in evaluating complex machine learning models, we recommend not only running the model deterministically (with any and all sources of randomness fixed to specific seeds), but also running the pipeline on multiple machines to determine whether this affects performance. We have demonstrated that this difference can vary unpredictably from one task to the next. Alternatively, those who are willing to disclaim exact replicability can run a model multiple times stochastically (some will test multiple “temperatures” of randomness(62)) to determine whether the overall findings are reproducible. However, any claim that the output is frozen and replicable must be backed by an additional run on a different CPU.

As it currently stands, we have demonstrated that public RNA-Seq data, while noisy, are still sufficient to prioritize candidate drug-gene relations, and when added to one’s repertoire of predictive tools, can increase the likelihood of finding a promising regulator without notably raising the risk of a false positive. Furthermore, as more new perturbations are tested, the recall of SNACKKSS will improve quadratically, and it could ultimately surpass the literature-based tools. Regardless of whether it does, we anticipate that RNA-and literature-based tools will continue to complement each other, and be better used together than alone to find pharmacologic gene regulators that could be repurposed for Mendelian disorders.

## Data availability

Our processing pipeline is publicly available, and divided into four GitHub repositories. The first, https://github.com/coledeisseroth/SNACKKSS_NLP (Zenodo DOI: 10.5281/zenodo.23025037), contains the materials needed to train and evaluate the optimal cascade of BERT models for GEO study curation. The second, https://github.com/coledeisseroth/SNACKKSS (Zenodo DOI: 10.5281/zenodo.19075797), takes the eight trained BERT models generated from the prior repository, and runs them on all of the available study metadata in GEO, then feeds the output into our default relationship-prediction pipeline (alongside the mouse-oriented version, which uses completely different data to make predictions). The third, https://github.com/coledeisseroth/SNACKKSS_Eval (Zenodo DOI: 10.5281/zenodo.19075806), runs the formal comparisons that we present in this article, evaluating the predictive ability of different variations of SNACKKSS and CMap, as well as PARMESAN and PubTator3. It takes as input the experiment files (gene_sample_controls.txt and drug_sample_controls.txt) generated from running the SNACKKSS pipeline.

The final repository, https://github.com/coledeisseroth/SNACKKSS_Revision (Zenodo DOI: 10.5281/zenodo.23024995), runs additional analyses requested during the revision process, and imports the Docker images and some data files from the SNACKKSS_NLP and SNACKKSS_Eval repositories. Our Docker images are available upon reasonable request, and can be generated anew using the Dockerfile in each GitHub repository. The raw output of our latest SNACKKSS curation run, and the latest predictions made using those curated studies, are available at snackkss.nrihub.org.

## Generative AI usage

We used Microsoft Copilot to learn the basic functionality of Docker, but we wrote our own code. In the process of developing SNACKKSS-2C, any studies for which AI (specifically, ChatGPT) was consulted for clarification are marked with a “(G)” in the “Curated_by” column of Supplemental table 3. Anthropic’s Claude provided editorial suggestions, each of which we reviewed manually and either rejected, or integrated after re-wording. Generative AI was not used for anything else in this study. BERT models were used only for classification, not for text generation (the latter of which they are not designed for). For the web interface (separate from the study), generative AI (specifically, Gemini) was used to make a browser icon, and provided some assistance in webpage formatting.

## Acknowledgements

We thank Seon Young Kim for her guidance in the design of the web portal. We also thank Daryl Scott, Ryan Dhindsa, Jimmy Holder, and the members of the labs of Drs. Huda Zoghbi (particularly Rebecca Meyer-Schuman, Ashley Anderson, Yan Li, and Mason Tate) and Zhandong Liu (particularly Hyun-Hwan Jeong) for their feedback, advice, and support.

## Author Contributions Statement

Conceptualization: C.A.D., Z.L, H.Y.Z.; Data curation: C.A.D., B.B., B.S.S.; Formal analysis: C.A.D.; Funding acquisition: C.A.D., Z.L., H.Y.Z.; Investigation: C.A.D.; Methodology: C.A.D., Z.L., H.Y.Z.; Project administration: C.A.D., Z.L., H.Y.Z.; Resources: Z.L.; Software: C.A.D., Z.S.; Supervision: C.A.D., Z.L., H.Y.Z.; Validation: C.A.D.; Visualization: C.A.D.; Writing – original draft: C.A.D.; Writing – review & editing: C.A.D., Z.L., H.Y.Z.

## Funding

This work was supported by the Eunice Kennedy Shriver National Institute of Child Health and Human Development [F30HD117505 to C.A.D.]; the National Institute on Aging [R01AG057339 to Z.L.]; the CHDI Foundation to Z.L.; the Huffington Foundation to Z.L.; the Chao Foundation to Z.L.; Howard Hughes Medical Institute to H.Y.Z.; and the National Institute of Neurological Disorders and Stroke [2R37NS027699 to H.Y.Z]. The content in this publication is solely the responsibility of the authors and does not necessarily represent the official views of the National Institutes of Health.

## Conflict of interest disclosure

C.A.D. has family who own equity in MapLight Therapeutics.

## Notes

https://github.com/coledeisseroth/SNACKKSS_NLP

https://github.com/coledeisseroth/SNACKKSS

https://github.com/coledeisseroth/SNACKKSS_Eval

https://github.com/coledeisseroth/SNACKKSS_Revision

### Summary of Updates

We fixed the model-saturation test and one of the Bonferroni multiple hypothesis corrections, and made editorial changes.

