## Supplemental methods, figures, and table legends for "Transcriptomic data and biomedical literature synergize in finding pharmacologic gene regulators"

**Supplemental materials**

**Supplemental methods**

**Training and testing GEO-curating BERT models**

For study classification, we only use the metadata fields “Title”, “Summary”, and “Overall design”. For sample and control classification, we use the “characteristics”, “description”, “organism”, “source_name”, “taxid”, and “title”. For target classification, we use those fields, as well as the protocol fields, as the rest of the sample description may fail to indicate what gene was being disrupted.

There were four fine-tuning regimens we tested for each BERT model: 1. training on CREEDS’ dataset alone; 2. training on our dataset, SNACKKSS-MC alone; 3. training on CREEDS’ dataset, and then training further on SNACKKSS-MC; 4. training on SNACKKSS-MC, and then training further on CREEDS’ dataset. GEO series IDs (GSE) from both CREEDS’ dataset and SNACKKSS-MC were shuffled with a random seed of “2025”, and split into three equal segments, and a smaller fourth one using Bash’s “split” command, which indexed divisions using the nomenclature “xaa, xab, xac, xad”. The cross-validation consisted of training a given model on three of the segments, and testing on the fourth. For any tests involving both SNACKKSS-MC and CREEDS’ dataset, we would maintain the same 4-fold architecture. In other words, when we tested the CREEDS-trained model on SNACKKSS-MC, the trained model being tested on SNACKKSS-MC’s “xaa” division would have seen neither SNACKKSS-MC’s nor CREEDS’ “xaa” division.

Because these BERT models cannot interpret more than 512 tokens at a time, we *chunk* larger bodies of text by breaking them in half until the segments are readable. As a safety buffer, we set the maximum token count to 500. For study, sample, and control classification, we assign a positive label if at least one of the chunked segments was labeled as positive.

BERT models assign each token (word fragment) in a sequence a vector-encoding based on the tokens that surround it(1). For target classification, they learn to classify each token as either the beginning token of a perturbagen, a non-beginning token thereof, or outside of any mention of a perturbagen (also known as the “BIO” format, Beginning, Inside, Outside(2)). All adjacent tokens labeled as positive are merged into one extracted entity name.

For study, sample, and control classification, we represent the question as a binary sequence classification task. BERT models add a special “[CLS]” token to the beginning of every sequence, and just like the other tokens, it receives a vector-encoding based on its neighbors. The model can be trained to classify this token alone, thus predicting what label should be assigned to it given what was observed in the text, and this approach is commonly used to classify the body of text as a whole. Training parameters are taken from the text- and token-classification tutorials from Hugging Face(3, 4), and are changed as needed to ensure functionality and address specific scientific questions. We include comments on the changed parameters in the SNACKKSS_NLP GitHub repository, in the “text_classification_finetune.py” and “target_finetune.py” scripts.

Specifically, the learning rate of 2x10^-5^, the epoch count of 2, and the weight-decay of 0.01 are suggested by the Hugging Face tutorials, while the use of gradient-checkpointing is intended to save memory, and the use of a random seed and requiring the use of a CPU are intended to test replicability across devices. The default batch size is 16, but for sequence classification (study, sample, and control), we had set the batch size to 1 while debugging the training procedure, while the token classification (target) kept its batch size of 16. Due to this discrepancy in batch size, our saturation test displayed in Supplemental table 7 and Supplemental figure 3 evaluates progressive training using both batch sizes, 1 and 16, for all tasks.

For training and testing our control classifier, we only classify pairs of samples if the manually curated dataset posits that the two samples differ by exactly one perturbation of a given type (gene-disruption or drug), which one has and the other lacks. Because BERT models are designed to classify tokens (and, by extension, bodies of text) rather than compare them, we must introduce pairs of sample descriptions in a format that the model can readily interpret. For a given putative control sample C for a known disrupted sample D, we first align each field in C’s description to its most similar field in D’s. We then create a string consisting of the field label (e.g. “genotype:”) followed by a concatenation of all of the non-aligned text in C’s description. This allows the BERT model to pick up relevant labels such as “NT”, “DMSO”, or “vehicle” in C, while ignoring labels that are identical to those in D. We concatenate all of these strings of differences across the different fields, and then chunk and classify this concatenation as described above.

For accuracy testing, we use the following metrics: For gene-disruption study classification (the drug-study curation is evaluated the same way, but we describe these metrics in the context of gene disruptions), a true positive is a GEO series that was correctly labeled as testing a gene disruption; a false positive is a series that does not test a gene disruption but was labeled as testing one; and a false negative is a series that tests a gene disruption, but was labeled as not testing one. For gene-disruption sample classification, a true positive is a GEO sample that was correctly labeled as having a gene disrupted; a false positive is a non-gene-disrupted sample marked as having a gene disrupted; and a false negative is a gene-disrupted sample labeled as not gene-disrupted. For gene target classification, a true positive is a character index range within the sample description that indicates the disrupted gene, which was perfectly identified by the AI (in other words, it captured the complete term and nothing outside of it); a false positive is an extracted index range that does not perfectly match a range describing a targeted gene (even if it overlaps); and a false negative is an index range describing a targeted gene, which was not perfectly extracted by the AI (hence, a non-perfect overlap would constitute both a false positive and a false negative).

For gene-disruption control classification, all evaluated pairs of samples involve one with a gene disruption that the other lacks, with their sets of disrupted genes being otherwise identical; a true positive is a pair that was correctly classified as a control-disrupted pair; a false positive is a pair that was falsely classified as control-disrupted; and a false negative is a true control-disrupted pair that was not classified as such. For control classification specifically, when counting the true positives, false positives, and false negatives, we downweigh them based on the number of potential controls evaluated for each sample. In other words, if one sample has 10 potential controls, each pair classified will add 0.1 to its corresponding count. This is because in our pipeline, thousands of accurate controls ascribed to one perturbed sample will only result in us properly interpreting one perturbed sample.

This training and evaluation pipeline was run twice on an Intel Xeon Silver 4114 central processing unit (CPU) (the output we use) to test replicability within the hardware, and once on an Intel Xeon Silver 4314 CPU to test the variability between hardware. To determine whether devices of identical architecture could produce identical output, we ran a portion of the SNACKKSS_NLP pipeline on another Intel Xeon Silver 4314 CPU, with the same Docker image. Specifically, we tested DistilBERT on the first division (labeled “xaa”) of SNACKKSS-MC after training it on the other three divisions of SNACKKSS-MC. The finalized run on the GEO metadata and downstream analyses were only run once (the pipelines under the repositories “SNACKKSS”, “SNACKKSS_Revision”, and “SNACKKSS_Eval”, on the same Intel Xeon Silver 4114, a different 4314 and another 4314, respectively). Re-running them would not be worth the processor-hours, since we already know that the exact output will not be replicable.

**Model saturation**

An NLP model is never truly saturated with training data, as there are infinitely many ways to describe an experiment, and the model can always benefit from more such examples. We can, however, still measure the relative effectiveness of new examples as we continue to add them, assessing for growth and diminishing returns. Growth is defined as a significant positive Pearson correlation between the relative progress through the epoch and the unsmoothed F1 score. In other words, the F1 score is increasing with the number of samples trained on. A lack of growth does not mean that the model was not learning to accurately classify studies (this is measured by comparing the final precision achieved to that of a random classifier); only that its precision-recall balance was not steadily improving as it progressed through the training data. Meanwhile, we define diminishing returns as a significant negative Pearson correlation between the training progress through the epoch and the change in F1 score relative to the previous timepoint (for example, after training on 20% versus on 10% of the samples). In other words, it measures whether the gains in the F1 score are declining as the model trains on more samples.

**Comparing the BERT models’ performance to random guessing**

To statistically compare the unsmoothed precision and recall of the BERT pipeline to that achieved by random guessing, we assumed that a baseline classifier would randomly guess positive or negative (with a 50% chance of each) for any study, sample, word, or putative sample-control pair. For study, sample, and control classification, this would have an expected recall of exactly 50%, while the expected precision would be the fraction of total classifiable cases that are positive per the manual annotation. This facilitates a binomial statistical test.

For control classification, we run the binomial test with and without the aforementioned sample-level weighting (where we downweigh sample-pairs belonging to a perturbed sample with numerous potential controls). While sample-level weighting better reflects the classifiers’ performance across the experiments they curate, the binomial test is more closely suited to unweighted measurements.

For the baseline target classifier, we simplified the calculation in a generous manner, such that the task is inherently easier for the baseline than it is for the BERT models (thus making it more difficult for the BERT pipeline to win). We assumed that the classifier would target whole words rather than tokens, with the target always being separated out from any larger words that it might be enclosed in. Thus, an “extracted entity” is a set of consecutive words that are collectively labeled as one entity, and a “valid entity” is a set of consecutive words that actually correspond to an entity of interest (drug or disrupted gene). The expected unsmoothed precision is the likelihood that any sequence of words labeled as an entity happens to be a valid one. This is calculated by adding together the likelihoods of all valid entities and dividing this sum by the sum of the likelihoods of all extracted entities. The likelihood of an extracted entity is the product of the likelihoods of each enclosed word being marked as positive and any surrounding words being marked as negative—thus, a product of multiple 0.5’s. Likewise, the expected recall is the average of the likelihoods of each valid entity.

For separating out words in a passage, we use all of the chunked sample descriptions, with words labeled in the “BIO” format (beginning, inside, outside) for named entity recognition. Since these are training corpora for cross-validation, everything will be triple-counted, but that does not affect the unsmoothed precision and recall estimates, because the numerator and denominator are both multiplied by three. The pipeline is available in the SNACKKSS_Revision repository.

Of note, we later found an error in our pipeline for processing the annotations, which missed 176 of the 1127 (15.6%) annotated experiments. This error did not introduce any fictitious experiments, and only caused a loss of training and testing cases, so we left the stable release as is, but we now make a note of this, both here and in the SNACKKSS_NLP repository.

**Accuracy testing**

In our original publication on PARMESAN(5), we measured the total number of regulatory relationships predicted at the smallest score achieving each directional accuracy (synonymous with what we call unsmoothed precision). The predictive measures we introduce in this study, however, are far more computationally intensive, requiring us to limit our evaluation to the regulatory relations that we can verify with DGIdb or Reactome. This gives us a measurement of recall, which is a less-informative measure of the absolute coverage of these tools, but is still effective for comparing different methods.

Additionally, in contrast to said publication, we have begun to eschew our reliance on the paid black-box statistical tools in GraphPad Prism, so that we could have our accuracy evaluations and hypothesis tests as a part of our open-source pipeline (or in Microsoft Excel, as needed). For evaluation of one predictor’s ability to prioritize correct relations over incorrect, we transitioned from Prism’s extra sum-of-squares F test (comparing the one-phase-decay constants of correct and incorrect as we increase the score threshold) to a log-rank test (still comparing the decay rates of correct and incorrect predictions as we increase the score threshold). For comparing different prediction tools, we transitioned from a Friedman rank-sum test measuring the number of predicted relations at integer-percent accuracy thresholds to a binomial test for the proportion of predictor A’s achieved unsmoothed precision-recall states where predictor B achieved simultaneously superior precision and recall. Granted, if a user knows what precision they want, then a statistical test that measures superiority across precision-recall states will not be helpful.

Furthermore, because we are testing many hypotheses, we apply Bonferroni correction to many of our p-values. For the binomial tests measuring whether SNACKKSS’ and CMap’s gene-perturbation samples show the expected expression changes in the target gene (**Supplemental table 9**) any of the tests returning positive would suggest that there are valid gene-perturbation signatures being identified.

When assessing the signature-based pipelines, and variations of each, for their ability to prioritize correct over incorrect (**Supplemental table 10**), any of them succeeding would speak in favor of signature-based relationship predictions. Importantly, we only account for the log-rank p-values from the leave-one-out cross-validation (LOOCV), not the ones at each of the ten thresholds, because we would not consider it a success if one of the pre-set thresholds yielded a strong prioritization ability. Likewise, when testing the non-signature-based predictors (**Table 2**), any of them succeeding would attest to the predictive ability of non-signature-based tools.

In our ablation test determining whether any predictive tool broadly benefits the precision and recall of the ensemble (**Supplemental table 16**), any tool succeeding would mean that there is a useful contributor to this effort that is worth our attention. Finally, when comparing the different signature-matching formulae (**Supplemental table 19**), this was a separate test setting from our other prioritization tests, answering the question of whether any of 3 signature-matching formulae effectively prioritize correct predictions over incorrect ones. Any of these log-rank tests returning positive would point to the validity of signature-matching in this unique setting.

Our pipeline for assigning unique Entrez IDs and PubChem Substance IDs (SIDs) to the relations in DGIdb misprocessed three of its relations. Two involved “HYODEOXYCHOLIC_ACID”, and the third involved “TETRALIN_UREA ANALOGUE (70)”. The resulting artifacts were fully contained, and did not introduce any fictitious relations that could have affected our accuracy evaluations—only the loss of three relations out of tens of thousands. Therefore, we are leaving the stable public release of SNACKKSS as is, but we make a note of this both here and in the SNACKKSS_Eval GitHub repository.

**CREEDS’ data were incorporated into our evaluation framework**

CREEDS’ manually annotated study sets had several artifacts that led to formatting issues, so we had to manually clean them. The cleaned, tab-delimited tables are in our GitHub repository, “SNACKKSS_NLP”. Additionally, we manually determined whether their descriptive study labels (i.e. not explicitly written as “KO” or “KD”) qualified as KO/KD studies. After making these adjustments, the data were suitable to feed into our training and testing pipeline, just as we did with SNACKKSS-MC.

**Synonym resolution**

When evaluating the models’ performance in target classification, we only consider whether the specific text describing the target was detected, not whether we were able to map the text to the correct gene product or drug. Unique identifiers for these entities will have overlapping sets of synonyms, making it difficult to establish a ground truth for benchmarking. Furthermore, we wanted to keep our NLP evaluation pipeline static by eschewing any reliance on the ever-changing lexica.

However, entity normalization(6) is necessary in large-scale meta-analyses, so for the finalized run of SNACKKSS on the GEO metadata, we link the gene targets to their Entrez IDs and the chemicals to their PubChem SIDs. Our normalization algorithm is as follows: we first find all potential IDs for each entity name extracted from the text. If one identified name shares an ID with a longer identified name, it will be assigned to whichever ID that longer name was assigned to. If an identified name has no extracted synonyms that are longer than itself, it is assigned to its numerically smallest ID.

**Filtering control classification data**

We use five criteria to limit the number of control classification instances. We describe them in the context of gene disruptions, but the same approach is taken for drug treatments. 1. The potential control must have all of the automatically identified gene disruptions in the perturbed sample except one. We handle this criterion differently between the accuracy testing and the finalized run: in the former, we index entities by their names alone, to minimize the susceptibility of our analysis to changing external databases. In the finalized run, however, which is already highly susceptible to this, we index genes by their Entrez IDs and chemicals by their PubChem SIDs. 2. The potential control must not have any automatically identified disruptions not found in the perturbed sample (except DMSO for drug studies, as it is a common vehicle that is often not mentioned in drug-treated sample descriptions). The same differences from Criterion 1 apply here. 3. The potential control must not differ from the perturbed sample in any of the fields that, in our manually curated dataset, we used to separate experiments (barring those labeled with “genotype”, “condition”, or “vector” for gene disruptions, and “exposure” or “treatment” for drugs). This filter does not apply to the accuracy tests, because that would compromise our evaluation. 4. The potential control must not differ from the perturbed sample in more than eight different description fields, which was the highest number of differing fields observed in a valid control in our manually curated dataset. This filter does not apply to the accuracy tests. 5. The GEO series containing these samples must have fewer than 10,000 comparisons to be made. This filter is solely for feasibility, as some studies can generate millions of sample pairs that need to be compared.

We run the top-performing control classifier on the first chunk of the alignment of each pair in this filtered list. Thus, for each target gene, we have a set of samples that are classified as having that gene disrupted, and for each of those samples, we have a set of appropriate control samples that do not have that specific gene disrupted. A sample with multiple gene disruptions can have multiple sets of controls, one for each disruption that it received.

**Handling samples from multiple read count datasets**

ARCHS4 collapses replicates into one GEO sample (one “GSM” identifier), whereas Recount3 and DEE2 do not collapse the replicates, and index their samples using their sequence read archive (SRA) run IDs (with “SRR” identifiers). If one perturbed sample has multiple runs, we treat each run as a separate sample with the same features. Likewise, if a control sample has multiple runs, all of its runs are taken as control samples.

All read counts are first converted to transcripts per million (TPM). For a given experimental sample with a disruption of gene A and at least two control samples, we use its controls to establish a mean and standard deviation for the TPM of each gene B. We would then calculate a z-score for the TPM of B in this disrupted sample. If a given sample has multiple disruptions, each with its own respective controls, we interpret this as two perturbed samples with different disruptions and normalize them accordingly. In other words, if Sample A is treated with Drugs X and Y, Samples B and C are treated with X, and Samples D and E are treated with Y; then we create two normalized samples from A: one treated with X and normalized to D and E; and one treated with Y and normalized to B and C.

We keep the three pre-computed read-count datasets completely separate from one another until we are calculating the consensus signature of a given gene’s disruption. When normalizing individual perturbed samples to their controls, we only use controls that are present in the same read count dataset. In other words, if Recount3 and DEE2 both have perturbed sample A, but Recount3 has controls B and C while DEE2 has controls C and D, we interpret this as two separate perturbed samples: Recount3’s Sample A (normalized to Recount3’s B and C) and DEE2’s Sample A (normalized to DEE2’s C and D).

**Connectivity Map processing**

We accessed the Connectivity Map (CMap), and downloaded its level-5 analysis. Because the data files are given in a unique binary format with the extension “.gctx” that takes a considerable amount of memory to open (the level-5 compound file took us approximately 34 gigabytes of memory), we first converted them into gzipped Unicode text files. The siginfo_beta.txt file was used to determine which collapsed, normalized samples were receiving gene-targeting shRNAs or being treated with small molecules. For a given perturbed sample, its controls had to be in the same cell line and have the same incubation time. Dosage was not taken into consideration—this proof-of-concept, if viable, should be robust to such a variable. For drug-treated samples, the controls had to have received a DMSO vehicle, and for shRNA-knockdown and CRISPR-knockout samples, any sample with the label “ctl_vector” was accepted as a control—if this method is viable, it should be robust to any variation among control vectors.

For relationship-identification purposes, our handling of CMap’s chemical and gene identifiers is slightly different from that for SNACKKSS. When calculating consensus signatures (nested Z-scores), we consider two chemical names distinct even if they likely refer to the same chemical. As an example, CMap uses compounds that it refers to as “C-646” and “C646”, which we assume to be the same (PubChem SID 85332784), but we consider them separately throughout the signature calculation and matching process. When testing accuracy and running LOOCV, we have to establish a one-to-one name-ID map, as this ensures a fair comparison to other tools and minimizes the risk of data leakage for the LOOCV test. When we do the conversion, we assign a single identifier to each named entity, and if multiple names used in CMap’s lexicon map to the same identifier, we only use the samples using the name that is found more often. We found 273 samples using “C-646” and 20 using “C646”, and DGIdb refers to the chemical as “C646”. We thus convert “C-646” to its PCSID, and ignore the “C646” experiments.

**Precision-recall curves**

To plot precision-recall curves for regulatory relationship predictors, we use Reactome’s functional interaction dataset(7) and DGIdb(8) as gold-standards for gene-gene and drug-gene relations (respectively). We consider supportive and inhibitory relations separately, and will use inhibitory drug-gene relations as an example. For a given predictor, we define a true positive as an inhibitory drug-gene relationship from DGIdb that the predictor identified as such, a false positive as a supportive drug-gene relationship that the predictor marked as inhibitory, and a false negative as an inhibitory drug-gene relationship that the predictor either did not identify or marked as supportive. Smoothed and unsmoothed precision and recall are then calculated as described in the Methods.

**Comparison of DF1 to the Spearman correlation**

The first task in comparing the performance of DF1 and the Spearman correlation is to evaluate SNACKKSS’ and SA4’s predictive ability at a given differentially expressed gene (DEG) threshold. To do this, we must correlate the signature of every gene disruption registered in SNACKKSS (2,606) with that of every drug registered by both SNACKKSS and DGIdb (492), and also with that of every gene registered by both SNACKKSS and Reactome (1,139). This is 1,139*2,605=2,967,095 correlations between gene-disruption signatures, and 492*2,606=1,282,152 gene-disruption/drug signature correlations, with a total of 4,249,247 matches to calculate. On a 64-core CPU, running up to 11 Spearman correlations simultaneously in Python (one correlation calculation can occupy multiple cores), we could calculate approximately 486 correlations per minute. To test one DEG threshold, it would thus take approximately 6 days, and testing 10 thresholds would take 61 days.

In contrast, we could calculate approximately 1,832 DF1 matches per minute, also running up to 11 jobs simultaneously. Testing one threshold with 11 cores would take approximately a day and a half, and testing 10 thresholds simultaneously can occupy 10 cores for approximately 16 days. Since each DF1 job (unlike the Spearman correlation function) only occupies one core, this method offers more capacity to parallelize, and allows us to test all of the variations that we did. When, for instance, running DF1 across 31 cores, we could calculate approximately 10,261 matches per minute.

For generating a resource (as opposed to benchmarking the tool against a gold-standard dataset), the computational load intensifies. We have 2,606 gene disruptions and 1,934 drugs tested on human tissue, and 3,103 gene disruptions and 1,334 drugs tested on mouse tissue. Having used a gold-standard dataset to decide on the optimal DEG threshold for four predictive tasks (supportive and inhibitory gene-gene and drug-gene relations), we will have to run this process four times. This would mean there are (2,606*1,934) + (2,606*2,605) + (3,103*3,102) + (3,103*1,334) = 5,040,004 + 6,788,630 + 9,625,506 + 4,139,402 = 25,593,542 matches to calculate per threshold, and 102,374,168 matches total. Running DF1 on 31 cores can take 7 days, while using the Spearman correlation 11 jobs at a time would take 146 days. This is why we use DF1 to match perturbation signatures at such a scale.

For benchmarking, due to the runtime of the Spearman formula, we had to streamline our LOOCV pipeline. The default approach involves predicting the regulatory relationships in Reactome and DGIdb for both SNACKKSS and SA4, at all tested DEG thresholds. Again, the Spearman correlations would occupy a 64-core CPU for 61 days if we did this. Instead, we use LOOCV to optimize the DEG threshold for predictive accuracy in SNACKKSS’ direct signature-matching (excluding all relations involving the left-out modulator), and then use only the optimized threshold to predict SA4’s targets for the left-out modulator. This reduces the runtime by 80%, and still ensures that we are not testing the pipeline on any data used to optimize the threshold.

Since running a Spearman correlation provides a p-value in addition to an effect size, we tested its predictive ability without p-value filtering, and requiring p < 0.05.

**Hardware requirements**

In order to re-run our pipelines, users will need approximately four terabytes of disk space (187, 2,656, 815, and 169 gigabytes to run the SNACKKSS_NLP, SNACKKSS, SNACKKSS_Eval, and SNACKKSS_Revision pipelines, respectively). We ran these pipelines on shared servers that were simultaneously running other jobs, so we cannot provide definitive runtime statistics. However, we can attest that the full process will span multiple months on our CPUs, so we recommend that users run it in the background on a machine that can simultaneously handle their daily computational needs.

The single highest-memory task in this study is the extraction of the Connectivity Map’s .gctx files, which requires up to 34 gigabytes of memory when running SNACKKSS_Eval. For the other three pipelines (SNACKKSS_NLP, SNACKKSS, and SNACKKSS_Revision), the highest-memory task is to run BioBERT or BioMedBERT, which requires approximately 15 gigabytes of memory.

**Supplemental tables**

**Supplemental table 1: Manual study annotations**

We manually annotated 625 GEO series. The columns are as follows:

Dataset ID: The series identifier, beginning with 200.

Accession: A redundant column, the series identifier beginning with “GSE”.

Curated_by: The first name of the individual who annotated the study.

Overexpression(OE)_Knockout(KO)_Knockdown(KD)_Other-gene-modulation(OM)_Drug(D)_None(N)_Gene-and-Drug(GD): The type of experiment being done. Each distinct controlled experiment within a series gets its own row. Studies that administer a perturbation (such as “KO”) but do not have any control samples for it may be labeled with “KO” or “N”, per the annotator’s discretion. While this does affect the training and testing of the study classifier, it does not do so in a way that conceptually goes against our training goals—the pipeline is designed to output no experiments from an “N” study or a “KO” study with no controls.

Sample_annotation(G=sample_id,T=Term): How we indicate control and perturbed samples in the following two columns. “G” means we list the sample accessions (GSM), separated by semicolons. T means that any samples with the indicated string in their descriptions are considered a part of that group; strings that have to both be present are separated by ampersands, and multiple strings that can indicate a sample without one another are separated by semicolons.

Control_samples: Our indication of which samples belong to the control group.

Perturbed_samples: Our indication of which samples belong to the perturbed group.

Comparison_Requirements: If this column is not empty, there are multiple experiments meeting the indicated criteria that must be considered separately, and are distinguished from one another by the field indicated (e.g. cell line). If multiple such fields are at play, they are separated by semicolons.

Input_Term: This column can be disregarded; it is a relic of an attempt to capture ChIP-Seq studies. We later concluded that our annotation schema is not sufficient to capture all types of high-throughput sequencing studies, and since we are currently only interested in RNA-Seq, we halted the effort to annotate other types.

Testing_Condition: String that must be present in the sample descriptions of both the control and the perturbed samples.

Perturbagen_terms: All strings (semicolon-separated) indicating the name of the perturbed gene or the chemical being administered. If multiple different perturbations are simultaneously occurring and are not isolated from each other, then their sets of terms can be separated with an ampersand (“&”).

Comments: Additional clarifications, written in plain language.

**Supplemental table 2: Corrected manual study annotations**

The modified study annotations, which we use for the pipeline. They are in the same format as Supplemental table 1. Note that the experiments previously classified as “GD” are now separated into individual experiments when possible.

**Supplemental table 3: Manual study annotation form a third individual**

An additional human curator (B.S.S.) manually annotated the same 615 studies that B.B. did, leading to the dataset that we call “SNACKKSS-2C”. Because this task was as much for educational purposes as scientific ones, the student was encouraged to consult C.A.D. for advice. The columns where C.A.D. was consulted were labeled with “(Q)” or “(q)” to indicate that these annotations were influenced by a prior curator. They were labeled with “(G)” if AI (specifically, ChatGPT) was consulted for clarification. The outdated “Dataset ID” and “Input_Term” columns are not present in this table.

**Supplemental table 4: Agreement between manual curators**

For binary study, sample, and control labels, we provide the number of labels that SNACKKSS-MC and SNACKKSS-2C agreed upon (“Agreed”), the number of labels in total (“Total”), the unsmoothed rate of agreement (“Observed agreement”), the expected rate of agreement if the two datasets labeled randomly at the rates that they did (“Expected agreement”), the Cohen’s Kappa value (“Kappa”), the binomial p-value for the rate of agreement (“Binomial p-value”), and the binomial 95% confidence interval around their rate of agreement (“Minimum” and “Maximum agreement”). Samples are only considered if both datasets agreed on the corresponding study label—in other words, we only observe whether they agreed on a gene-disruption sample if they agreed that that sample’s study was testing a gene-disruption. Likewise, we only observe target and control classifications for a sample if both datasets agree that that sample received the perturbation in question. Target classifications had to be handled differently, because rather than binary labels, they are extracted strings of text, and p-values could not be calculated to assess for non-random agreement because a theoretical random curator would pull random excerpts that would be infinitesimally likely to match those from another curator. We instead provide the labels that the two datasets shared (“Shared”), and those that were only in SNACKKSS-MC (“SNACKKSS-MC only”) and SNACKKSS-2C (“SNACKKSS-2C only”). We calculate the datasets’ unsmoothed precision in detecting each other’s target labels, and an unsmoothed F1 score where one of the two precision values is treated as recall (it does not matter which one). We then calculate 95% binomial confidence intervals around said precision values and F1 scores. For control classification, we statistically compare the two datasets with and without sample-level weighting (downweighing numerous pairs that all pertain to the same perturbed sample), as the weighting better captures the classification performance across samples, but the unweighted values are a closer fit for the binomial test.

**Supplemental table 5: Cross-validation split**

We used Bash’s “shuf” and “split” commands to establish our 4-fold cross-validation cohorts. In this table, we list the GEO series IDs belonging to each cohort, xaa-xad, for both SNACKKSS-MC, and CREEDS’ dataset.

**Supplemental table 6: GEO metadata NLP performance**

We provide the 4-fold cross-validation results from the NLP pipeline classifying GEO metadata. For each classification task, training each model on each manually curated corpus on each machine, we provide the number of true positives, false positives, and false negatives achieved on our SNACKKSS-MC dataset, and on that from CREEDS. For control classification, we downweigh the counts by the number of potential controls evaluated for a given disrupted sample. For example, if a disrupted sample has 10 candidate controls, each pair will count as 0.1 cases. We also provide smoothed and unsmoothed precision, recall, and F1 scores against SNACKKSS-MC, and binomial 95% confidence intervals (“minimum” to “maximum”) around the unsmoothed precision, recall, and F1 scores, and the absolute difference between the two devices’ unsmoothed F1 scores. This table supplies the values in Figure 1A.

**Supplemental table 7: Performance of BioMedBERT throughout its fine-tuning**

Running BioMedBERT on a 64-core CPU, training it on CREEDS’ dataset and then training that fine-tuned model on SNACKKSS-MC, we plot the smoothed and unsmoothed precision, recall, and F1 scores as it progresses through the training. We do this for study, sample, target, and control classification for both gene disruptions and drug treatments. For the first epoch of training on CREEDS’ dataset and the first epoch of training on SNACKKSS-MC, we save the model at a minimum of 10 equally spaced checkpoints. In order to calculate the steps prior to saving a checkpoint, we take the number of training samples in that cross-validation division, divide it by the batch size, divide that quotient by 11, and then round down to the integer. The progress value of 0 does not mean the model has not started training yet; rather, it means that the model has reached the first checkpoint of training, which covers less than 10% of the first epoch. This table supplies the values plotted in Supplemental figure 3. While BioMedBERT is being trained on CREEDS’ dataset, we also plot BioMedBERT’s testing performance on that same dataset. Additionally, we run this test using training batch sizes of both 1 and 16. For the first epoch of training, we measure the change in the unsmoothed F1 score at each timepoint, relative to the previous checkpoint or saved model (“Unsmoothed F1 score change since previous checkpoint”, provided there is an available previous score), and provide our statistical tests of continual growth (“Pearson correlation: Growth” and “Pearson p-value: Growth”) and diminishing returns (“Pearson correlation: Diminishing returns” and “Pearson p-value: Diminishing returns”) as the model progresses through the training data. The “Saturated?” column indicates whether the model’s training progress met our definition of model saturation.

**Supplemental table 8: Leave-one-out cross-validation for running BERT models on SNACKKSS-MC**

To compare the performance of the two CPUs after letting each one train, select, and use the most-accurate BERT models, we ran a form of leave-one-out cross-validation (LOOCV), where having trained each model on the other three 4-fold cross-validation (4FCV) splits and tested it on the fourth, we use all of the GEO series from the testing split except one to determine which model has the highest smoothed F1 score, and use that model’s predictions for that left-out series. For each classification task (study, sample, target, and control classification, for gene-disruption and drug studies, performed by a 40- or 64-core central processing unit (CPU), we provide the number of true positives, false positives, and false negatives. We also provide the smoothed and unsmoothed precision, recall, and F1 scores, and the 95% binomial confidence intervals around them (“minimum” and “maximum”, and the up- and down-error-bar sizes). For control classification, we run these analyses with and without sample-weighting, since the latter, despite being skewed by samples with a large number of potential controls, is more closely suited to a binomial test. Lastly, we provide the number of labeled cases in SNACKKS-MC that were manually classified as positive and negative (for target classification, this is calculated by summing the likelihoods of valid and extractable entities, as described in the Supplemental methods), the expected precision and recall achieved by random guessing, and the binomial p-values contrasting the BERT models’ performance from that of the random classifier. This table corresponds to Figure 1B.

**Supplemental table 9: Differential expression of targeted genes**

For gene-disrupted samples curated by SNACKKSS and the Connectivity Map (CMap), we provide the number of samples where the supposedly disrupted gene increased or decreased in expression, relative to its respective controls. We show this count at baseline, and as one requires a larger expression Z-score to consider a gene differentially expressed (“Minimum expression |z-score|”). For SNACKKSS, there are three databases (ARCHS4, Recount3, and DEE2), and each one has data from two species (Human and Mouse). For CMap, we separately consider samples whose supposedly targeted gene was among the “Landmark”, “Best-inferred”, or “Inferred” genes from their assay, and we also separate samples that underwent a “knockout”, “knockdown”, or “overexpression”. At the top, we provide the total number of samples in the measured group (for example, there were 19,025 gene-disrupted samples curated by SNACKKSS with data in ARCHS4), the overall proportion of these samples that showed decreased expression (“Proportion decreased”, 81.0%), the lower and upper bounds of the binomial 95% confidence interval around this percentage (80.4-81.6%), the binomial p-value for this proportion with an expected value of 50% (“Binomial p”, 0), and the Bonferroni-corrected p-value, where SNACKKSS and CMap have 6 and 9 hypotheses, respectively (“Binomial padj”, 0). This table corresponds to Supplemental figures 4 and 7.

**Supplemental table 10: Log-rank tests of DF1 signature-matching at different DEG z-score thresholds**

This table provides the log-rank test statistics for each SNACKKSS- (“SNACKKSS”) and CMap-derived (“ConnectivityMap”) predictor’s ability to prioritize correct relations over incorrect ones. We separately test their ability to predict gene-gene (“gene”) and drug-gene (“drug”) relations, and evaluate their prioritization ability overall (“all”) and when limiting to supportive (“pos”) and inhibitory (“neg“) relationship predictions, We measure the performance at each z-score threshold (0-0.9) above which one will accept a differentially expressed gene (DEG). Our main performance metric, however, is the performance achieved after deciding the threshold through leave-one-out cross-validation (LOOCV), and for this trial, we plot the log-rank test statistic (“LOO statistic”) and p-value (“LOO pvalue”). Since we are technically introducing 72 different relationship predictors, and the success of any of them could theoretically be considered an overall success, we run Bonferroni correction on the log-rank p-values, multiplying all of them by 72 (“LOO padj”, see the Statistical Analyses section in the Methods for a breakdown of the hypothesis count). In addition to SNACKKSS (“default”), we display the same performance metrics for all of its variations: requiring gene-disruption samples to have decreased expression of the supposed target (“targdown”), only using read count data from ARCHS4 (“archs4”), and using mouse data instead of human (“mouse”). For CMap, there is also a “default” setting, and the variations include limiting to shRNA knockdown data (“shrna_only”), including inferred gene expression levels (“inferred”), limiting to data taken from MCF7 cells (“mcf7”), and refining gene-disruption DEGs using overexpression data (“oe_corrected”). Additionally, we derive predictions from direct signature-matching (“f1_matches”), or from linking those signature matches to ARCHS4’s correlations (“archs4_correlation_linked”, i.e. SA4 and CMA4).

**Supplemental table 11: Correct and incorrect predictions from direct signature-matching-based predictors**

We display the numbers of correct and incorrect relationship predictions made by each signature-matching-based predictive tool, upon limiting the predictions to those within each score quantile. For example, in the quantile of 0.8, we only accept the top-20%-scoring predictions. For each predictor, we separately evaluate gene-gene (“Gene”) and drug-gene (“Drug”) regulatory relationships, using the manually curated databases (MCDBs) from Reactome and DGIdb (respectively) as gold standards. A “true supportive” or “false supportive” is a prediction of a supportive relation that, according to the MCDB, is supportive or inhibitory, respectively; likewise, a “true inhibitory” or “false inhibitory” is a relation predicted to be inhibitory, where the MCDB claimed it to be inhibitory or supportive, respectively. “SNACKKSS DF1” matches signatures curated by SNACKKSS, while “CMap DF1” matches consensus signatures from the Connectivity Map (CMap). In addition to the “default” pipelines for each one, we test the performance of each variation of the pipeline. For SNACKKSS, “ARCHS4-only” only uses read counts from ARCHS4, not from Recount3 or DEE2; “Mouse” uses mouse data instead of human, and “Target-down” only accepts gene-disruption samples if the supposedly targeted gene showed decreased expression relative to the controls. For CMap, “shRNA-only” ignores the CRISPR data, “MCF7” limits to samples from the MCF7 cell line, “OE-corrected” refines differentially expressed gene (DEG) lists using overexpression data, and “inferred” uses all inferred expression levels, rather than just the landmark genes.

**Supplemental table 12: Correct and incorrect counts for non-signature-based predictors**

This table has the same format as Supplemental table 11, but evaluates the performance of non-signature-based predictive tools. “PARMESAN consensus” and PubTator3 consensus” run PARMESAN’s consensus algorithm on relationships extracted by PARMESAN and PubTator3, respectively. “PARMESAN indirect” and “PubTator3 indirect” run PARMESAN’s indirect hypothesis formula on the relationships from PARMESAN and PubTator3, respectively. “PA4” and “P3A4” link the consensus relationships from PARMESAN and PubTator3 (respectively) to the correlations from ARCHS4 to make these predictions. “A4C” refers to predicting regulatory effects using just the coexpression matrices from ARCHS4, and we test their accuracy using either human (“Human”) or mouse (“Mouse”) data.

**Supplemental table 13: Correct and incorrect counts for predictors that link signature matches to ARCHS4’s correlations**

This table has the same format as Supplemental table 11, but evaluates predictions made by linking each signature-matching-based predictor to ARCHS4’s correlations. “SA4” and “CMA4” link the signature-matches from SNACKKSS and the Connectivity Map (respectively) to these correlations.

**Supplemental table 14: Justification of the top-scoring inhibitory drug-gene relation from SA4**

We provide the evidence supporting the top-scoring inhibitory drug-gene relationship prediction from the SNACKKSS/ARCHS4 hybrid (SA4). The drug would have a similar or opposite signature to an intermediate gene, whose expression would have a strong correlation with that of the target gene. The columns are the intermediate gene’s Entrez ID, the DF1 score from the drug to the intermediate, the expression correlation between the intermediate and target genes, and the score for this individual link, signifying its contribution to the overall conclusion.

**Supplemental table 15: Ablation test**

We use leave-one-out cross-validation (LOOCV) to have each predictive tool estimate its own accuracy, then believe the most confident predictor regarding an unseen regulatory relationship. We display the number of correct predictions made above the lowest confidence level that yielded the desired precision (Separately evaluating smoothed and unsmoothed precision). We test this performance using 10 different predictors (or 8 for drug-gene relations) in the “Nothing” column, and after removing each predictor from the cohort-- “PARMESAN consensus”, “PubTator3 consensus”, “PARMESAN indirect”, “PubTator3 indirect”, “PA4”, “P3A4” “Human A4C”, “Mouse A4C”, “CMA4”, and “SA4”. We separately evaluate supportive and inhibitory gene-gene and drug-gene relations. A valuable contribution from a predictor is signified by inferior performance after removing it—in other words, its column has smaller correct prediction counts (i.e. lower recall) than the “Nothing” column. For the purposes of calculating recall, we also provide the number of predictable relations in the gold-standard dataset, in the “Count” row. This table directly corresponds to Figure 4 and Supplemental figure 9.

**Supplemental table 16: Ablation test statistics**

For each predictor evaluated in the ablation test (Supplemental table 15), we statistically evaluate the improvement (or lack thereof) from adding it to our repertoire. In the case of SA4, for every distinct precision-recall state achieved using all predictors except SA4, we determine whether the combination of all predictors (including SA4) achieved a point of simultaneously higher precision and recall. The number of precision-recall states that improved with the inclusion of SA4 is provided under “Thresholds improved”, and the number that did not improve is under “Thresholds not improved”. Likewise, we calculate the proportion of these precision-recall states that improved (“Fraction improved”), after adding 1 to the denominator (“Smoothed fraction improved”), calculate a binomial p-value for whether the unsmoothed proportion is different from 50% (p-value), run Bonferroni correction on this p-value for 36 hypotheses (“Bonferroni-corrected p-value”, see the Statistical Analyses section in the Methods for a breakdown of the number of hypotheses), and calculate a binomial 95% confidence interval (smoothed and unsmoothed) around this proportion (“Lower-“ and “Upper bound fraction smoothed/unsmoothed improved”), and the highest smoothed and unsmoothed precision achieved under the given setup (“Maximum smoothed/unsmoothed precision”). We separately evaluate predictions of supportive and inhibitory gene-gene and drug-gene relations. The “Nothing” rows are a negative control, which should have zero improvements, since it will be equal to our baseline.

**Supplemental table 17: Leave-one-out cross-validation with each predictor alone**

In the same format as our ablation test (Supplemental table 15), we display the results of using each predictor alone (rather than using all predictors except one), where through leave-one-out cross-validation, we have the predictor estimate its own accuracy and make predictions on the masked relations. This supplies the “Champion alone” curves in Figure 4 and Supplemental figure 9.

**Supplemental table 18: Inhibitory drug target coverage with and without SA4**

Since SA4 shows a clear precision-recall benefit in identifying inhibitory drugs in DGIdb, even alongside other predictive tools, we measured the improvement in coverage in a different way. For any smoothed or unsmoothed precision one is willing to accept, we measure the number of genes for which an inhibitory drug was correctly identified, when using all predictors from Supplemental table 15 (“With SA4”), or using all of them except SA4 (“Without SA4”). We also display the difference between the two (“Delta”) and the “Fold-increase” for visualization purposes. This table directly corresponds to Supplemental Figure 10.

**Supplemental table 19: Predictive ability from using DF1 or Spearman correlations**

We test the ability of SNACKKSS’ signature-matching and SA4 to predict supportive and inhibitory gene-gene and drug-gene relations, matching perturbation signatures using either DF1, a Spearman correlation without a p-value requirement, or a Spearman correlation requiring a p-value below 0.05. At each level of precision (both smoothed and unsmoothed) in increments of 0.01, we show the number of correctly identified regulatory relationships achieved by each predictor. We also provide the log-rank p-values, uncorrected and Bonferroni-corrected for 24 hypotheses, testing each predictor’s ability to prioritize correct predictions over incorrect ones.

**Supplemental table 20: Performance of each predictor, using a new gold-standard dataset**

For each predictive tool—PARMESAN consensus, PARMESAN indirect, PA4, PubTator3 consensus, PubTator3 indirect, P3A4, Human and Mouse A4C, SNACKKSS, Connectivity Map, SA4, and CMA4—we measure the performance in predicting the relations in TRRUST v2 (gene-gene relations) and an amalgam of drug-gene relationship datasets (Drug Repurposing Hub, PharmGKB, ChEMBL, and Guide to Pharmacology), with any Reactome and DGIdb relations excluded. For each predictor, we provide the log-rank test statistic and p-value measuring its ability to prioritize correct predictions over incorrect ones (we do not run Bonferroni correction, because we are only interested in the performance of SA4 in predicting inhibitory drugs), and display the number of correctly identified relations when achieving each level of precision (smoothed and unsmoothed precision are plotted separately). We also provide in the “Count” row the number of identifiable relations in each new gold-standard dataset.

**Supplemental table 21: Ablation test, using a new gold standard**

In the same format as Supplemental table 15, we re-run the ablation test, using the entirety of Reactome and DGIdb to estimate precision (instead of leaving one modifier and its targets out), and using the aforementioned new gold-standard datasets used in Supplemental table 20. We also supply the number of predictable relations in the “Count” row, and the number of precision-recall states that were and were not strictly improved upon (better precision and recall simultaneously) upon adding the predictor in question (“Thresholds improved”, “Thresholds not improved”), and the binomial p-value comparing the fraction improved upon to 50%.

**Supplemental figures**

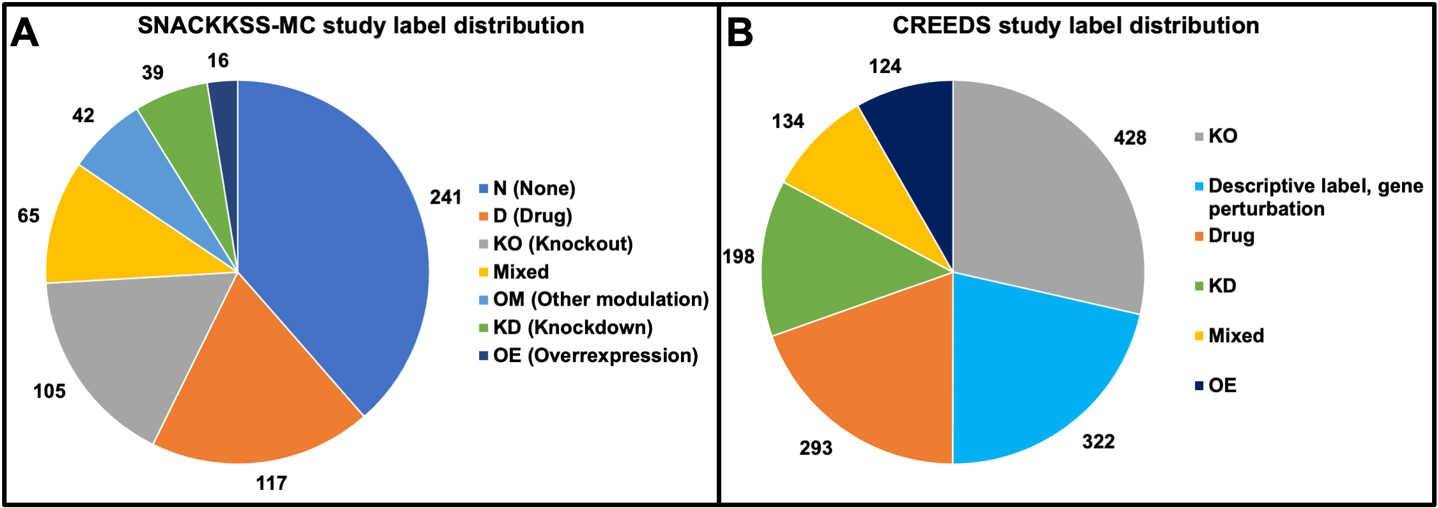

**Supplemental Figure 1: Types of GEO series observed in the manual curation**

We display the number of GEO series in our dataset, SNACKKSS-MC (**A**); and CREEDS’ dataset (**B**); labeled as containing each type of experiment. “Mixed” means that there were multiple experiment entries for this series, which did not all have the same label. “Descriptive label, gene perturbation” means that a given study from CREEDS’ single-gene cohort had a plain-text label other than “KO”, “KD”, or “OE”.

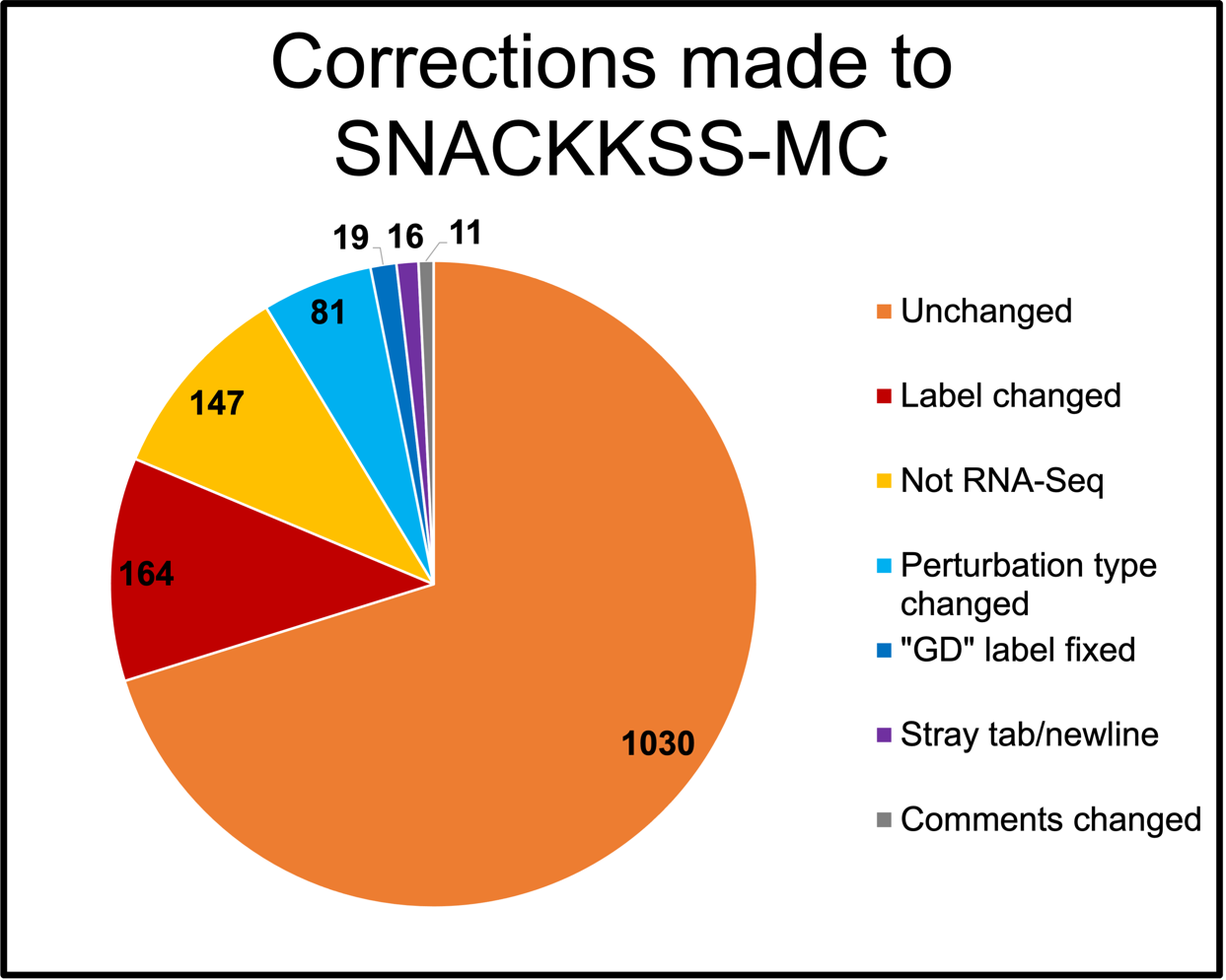

**Supplemental figure 2: Corrections made to the original SNACKKSS-MC dataset**

After SNACKKSS-MC was first curated by B.B., C.A.D. looked through the curated dataset and made corrections. Each line in the dataset corresponds to an individual experiment within a study, so we count the number of lines to which C.A.D. made each type of correction. “Unchanged” means no corrections were made. “Label changed” means one of the sample labels was corrected. “Not RNA-Seq” means that this experiment did not involve RNA-Sequencing samples, and its validity was not applicable. “Perturbation type changed” means the type of perturbation (Knockout, knockdown, overexpression, drug, other gene modulation, or nothing) was corrected. ‘”GD” label fixed’ means that the line was using an outdated perturbation type label, “GD”. “Stray tab/newline” means that extraneous whitespace characters were removed, because they interfered with the table’s formatting. “Comments changed” means that the final column, the comments section, was edited.

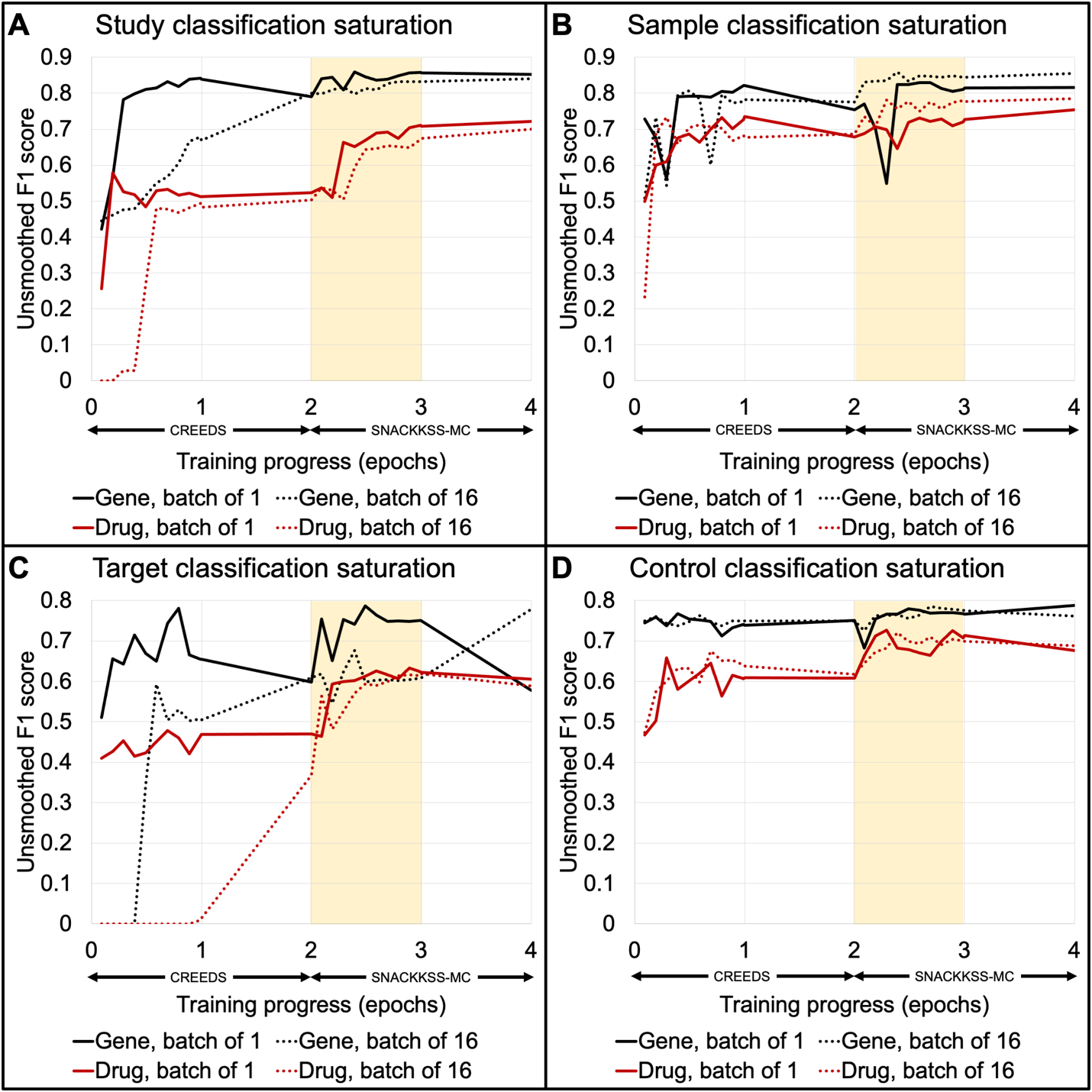

**Supplemental figure 3: Performance of BioMedBERT on SNACKKSS-MC throughout its fine-tuning**

We use 4-fold cross-validation to fine-tune BioMedBERT on CREEDS’ dataset (0<X≤2), and then further train that fine-tuned model on SNACKKSS-MC (2<X≤4), for study (**A**), sample (**B**), target (**C**), and control (**D**) classification. The X axis is the number of epochs trained on, the first two on CREEDS’ dataset and the third and fourth on SNACKKSS-MC. At X=0, the model has not yet begun training. At X=2, the model has finished two epochs of training on CREEDS’ dataset, and not begun training on SNACKKSS-MC. At X=4, the model has finished two epochs of training on SNACKKSS-MC, after having already had two epochs of training on CREEDS’ dataset. To distinguish the intermediate checkpoints from the finalized models, we increment the former’s X-values by 1/11—for example, at the first checkpoint of training on CREEDS’ dataset (progress = 0 in Supplemental table 7), X = 0.091. We highlight the section from 2<X≤3, which displays the progression through one epoch of training on SNACKKSS-MC, thus assessing the model’s saturation with the amount of training data it was given. The Y axis is the unsmoothed F1 score in predicting the annotations of SNACKKSS-MC. We plot this change in performance for each task—study, sample, target, and control classification for drugs and gene-disruptions. After finishing the training on CREEDS and then SNACKKSS-MC, two epochs each, the batch size did not change the final F1 score by more than 5%, except in gene-disruption target classification, for which there was a 20% difference. In this task, the batch size of 1 may have led to overfitting to the training data; while the batch size of 16 may have led the model to need the additional steps of optimization in the second epoch to finally recognize key patterns.

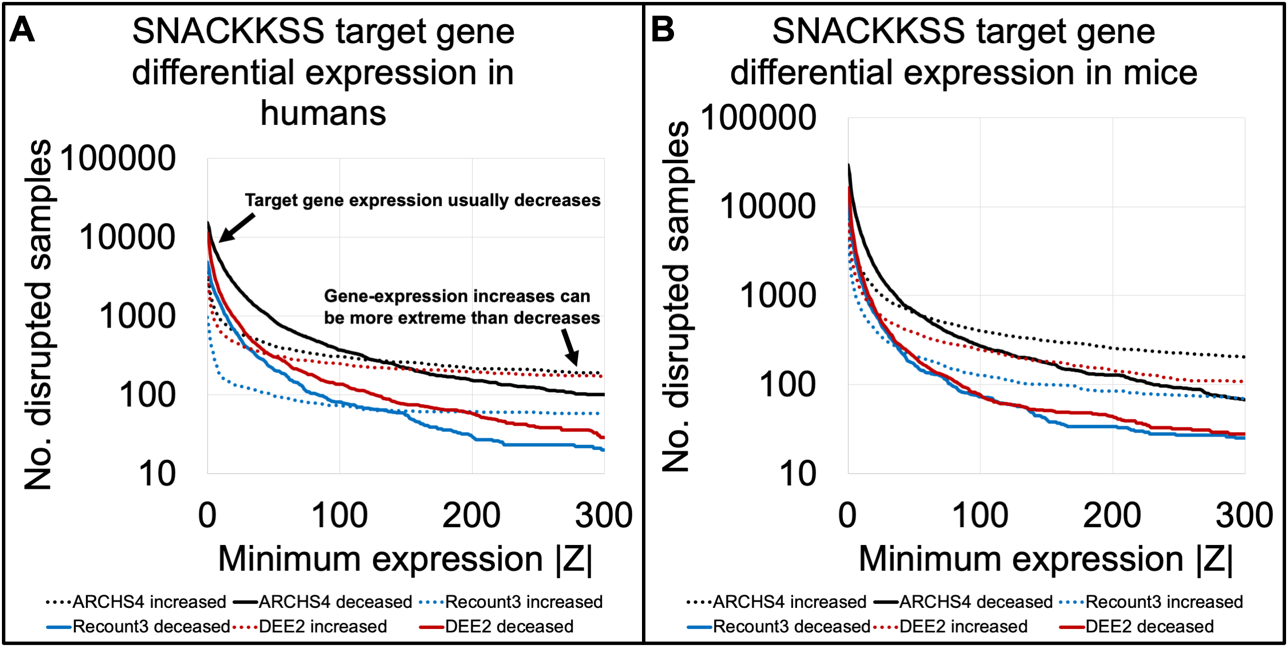

**Supplemental figure 4: Z-scores of the supposed target genes across normalized gene-disruption samples**

For all human (**A**) and mouse (**B**) read-count data taken from either ARCHS4, Recount3, or DEE2, we identify the gene-disrupted samples, and measure the z-score of the expression of the gene that was allegedly disrupted, relative to that sample’s controls (difference from the control mean divided by the standard deviation among the controls). The X axis is the minimum absolute z-score accepted, and the Y axis is the number of samples whose targeted gene had an absolute z-score above that threshold, with reduced (solid lines) or increased (dotted lines) expression of that target gene. As expected, the gene-disrupted samples predominantly showed decreased mRNA levels of their supposed KO/KD targets in all cases—except at the extremes, as the increases could be substantially stronger than the decreases. This is unsurprising, because read counts can increase to infinity, but cannot go below zero.

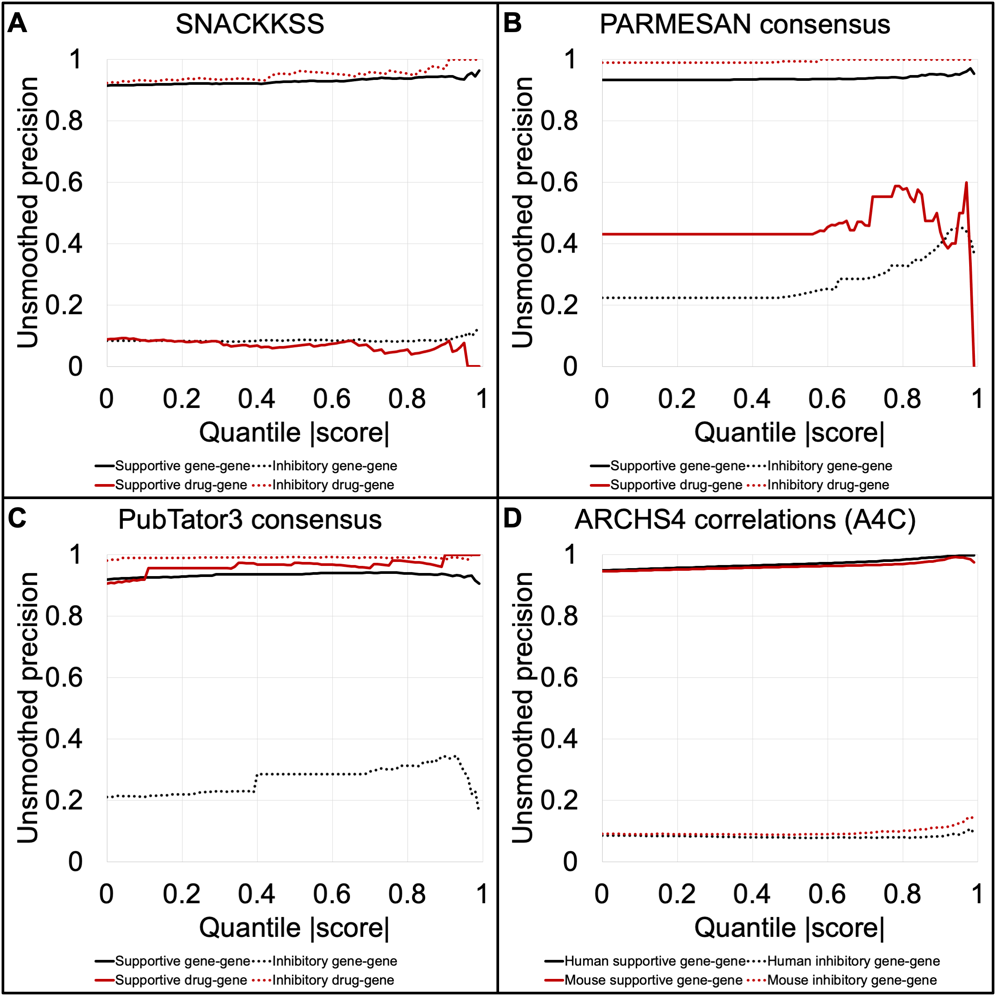

**Supplemental figure 5: Unsmoothed precision at increasing prediction scores**

This figure is formatted the same as Figure 2, except we measure unsmoothed precision instead of smoothed, for SNACKKSS’ signature-matches (**A**), PARMESAN and PubTator3’s consensuses (**B** and **C** respectively), and the gene expression correlations from ARCHS4 (**D**).

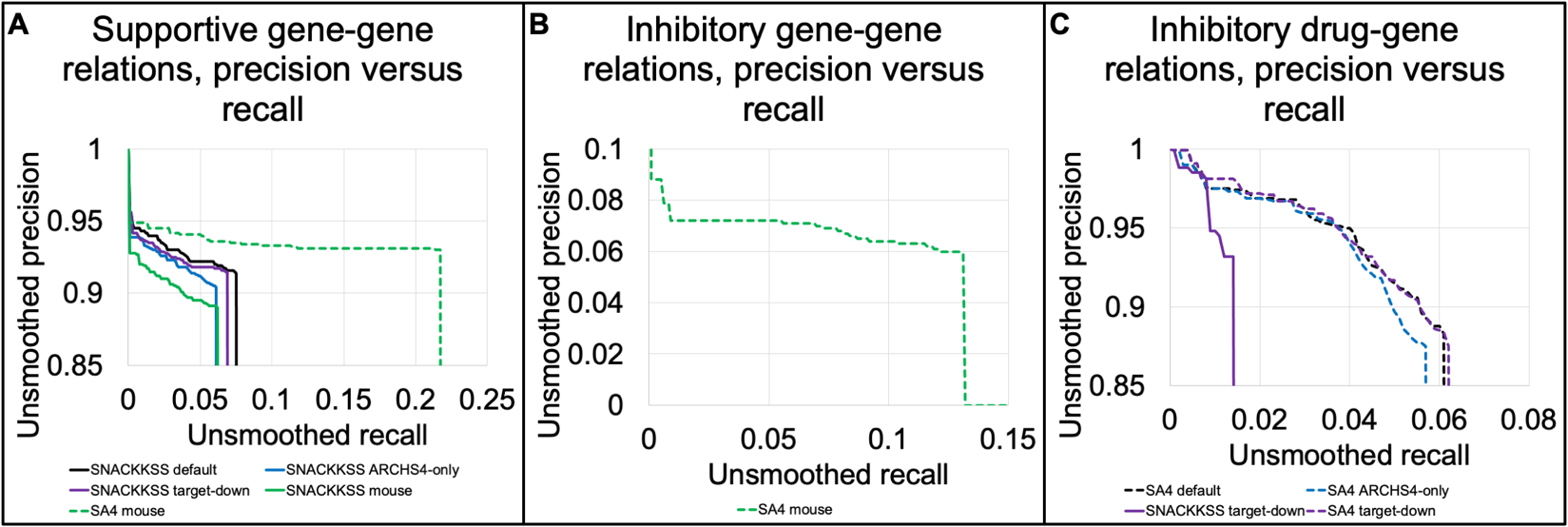

**Supplemental Figure 6: Evaluation of variations to SNACKKSS-based predictors**

Alongside the default signature-matching approach for SNACKKSS (“SNACKKSS”) and the predictions derived from linking them to ARCHS4’s correlations (“SA4”), we test three high-level variations to SNACKKSS and SA4 for their ability to prioritize correct regulatory relationships. The variations require decreased expression of the supposedly disrupted gene relative to the control samples (“target-down”), eschew the use of Recount3 and DEE2 (“ARCHS4-only”), and use mouse data instead of human (“mouse”). We separately compare their performance for supportive gene-gene (**A**), inhibitory gene-gene (**B**), and inhibitory drug-gene relations (**C**). We only plot a predictor if its correct predictions outlasted the incorrect ones with rising score thresholds, with an unadjusted log-rank p < 0.05. Since no setup effectively prioritized supportive drug-gene relations, we do not have a panel for this task.

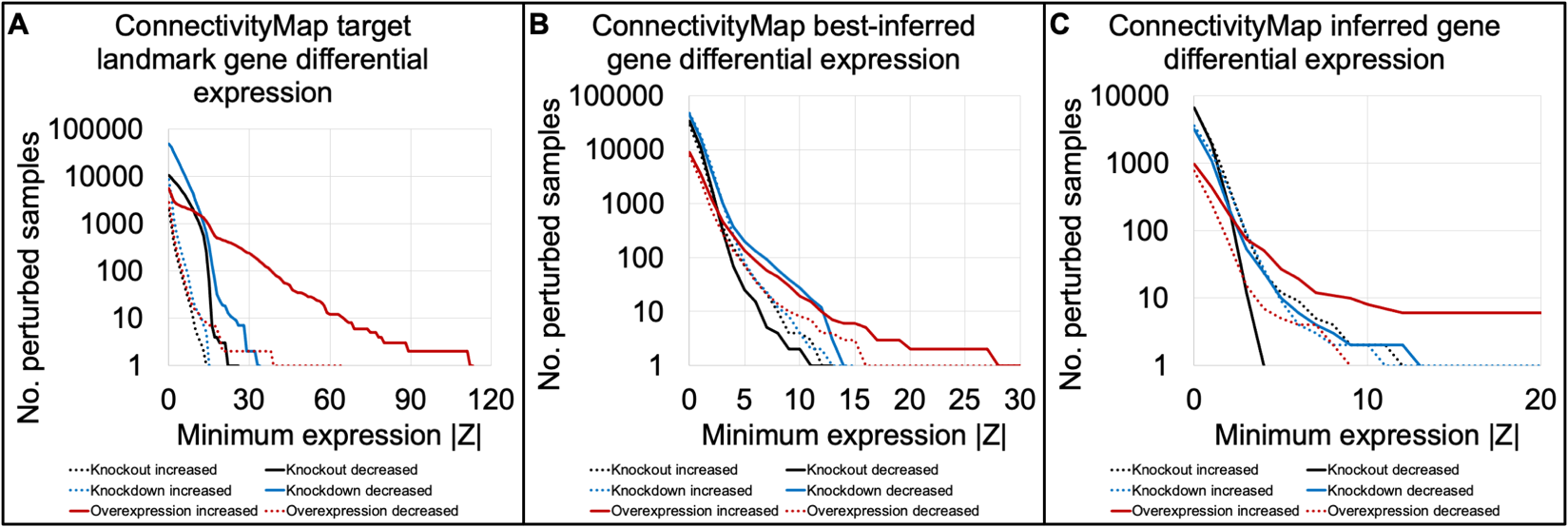

**Supplemental figure 7: Differential expression of supposedly targeted genes in the Connectivity Map**

This figure is formatted in the same way as Supplemental Figure 4, where we plot the number of gene-perturbed samples from the Connectivity Map (CMap) that, relative to their controls, had increased or decreased expression of their supposedly targeted gene. We plot this separately for shRNA-knockdown, CRISPR-knockout, and overexpression samples. Because CMap only directly measures the levels of 978 genes, we separately analyzed those “landmark” genes (**A**), the “best inferred” genes whose inferred levels supposedly correlated with actual expression levels (**B**), and the “inferred” genes whose inferred levels supposedly did not correlate with actual expression levels (**C**).

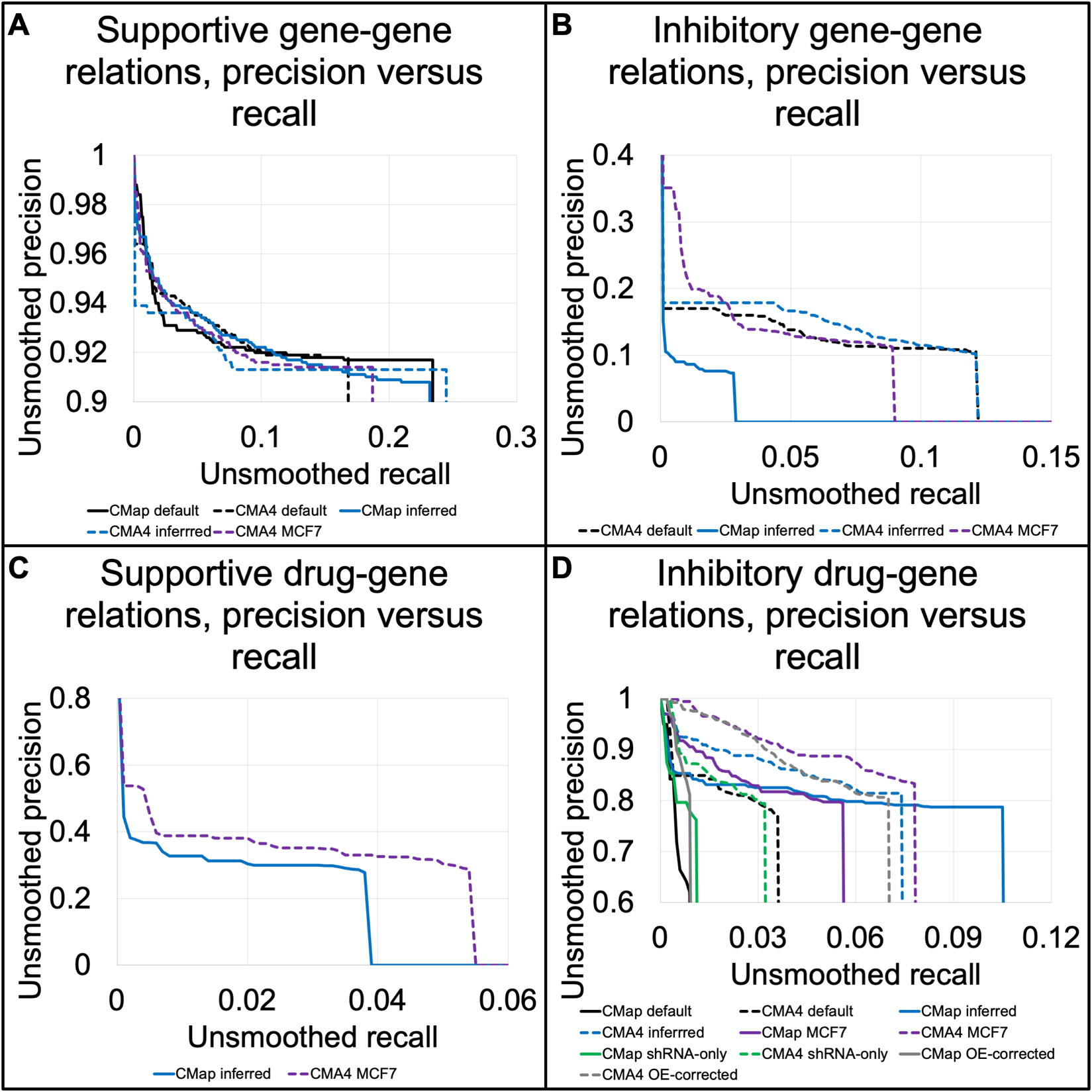

**Supplemental figure 8: Evaluation of variations to the Connectivity-Map-based predictors**

We display the performance of matching Connectivity Map signatures (“CMap”) and linking those matches to ARCHS4’s gene expression correlations (“CMA4”) using default parameters (“default”) or one of the four variations to the pipeline. Specifically, we limit the gene-disruption data to shRNA knockdowns (“shRNA-only”), include the inferred gene expression levels as part of the signatures (“inferred”), exclusively use data taken from MCF7 cells (“MCF7”), and limit the gene-disruption differentially expressed genes to those that went in the opposite direction upon overexpression of that gene (“OE-corrected”). We plot unsmoothed precision against recall in identifying, per manually curated databases (MCDBs), supportive gene-gene (**A**), inhibitory gene-gene (**B**), supportive drug-gene (**C**), and inhibitory drug-gene relations (**D**). We only display a predictor if its correct predictions outlasted the incorrect ones as the score threshold rose, with an uncorrected log-rank p < 0.05.

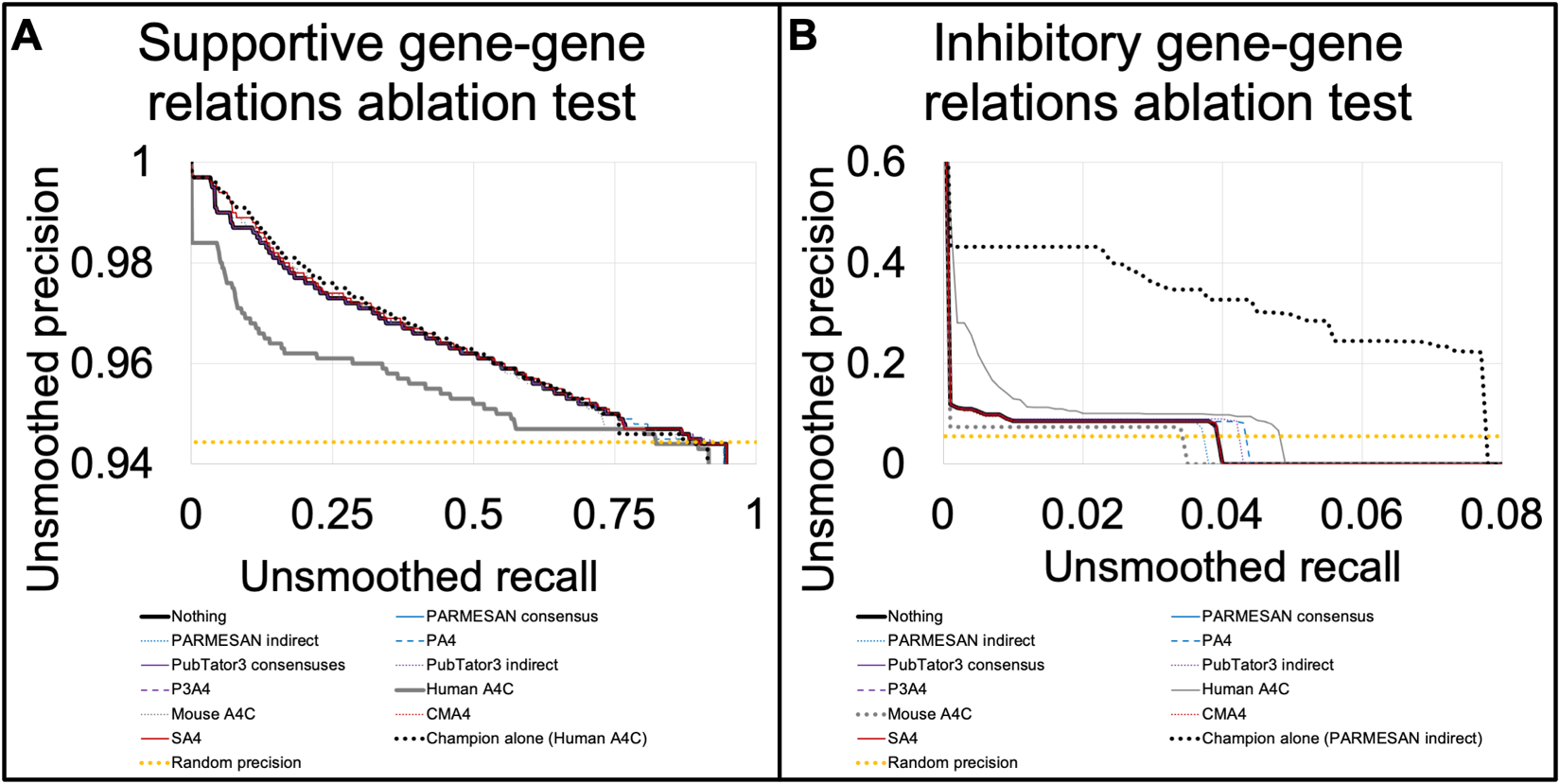

**Supplemental figure 9: Contribution of each predictor to identification of gene-gene relations**

In the same format as Figure 4, we test the performance of using all tested predictors in an ensemble to identify supportive (**A**) and inhibitory (**B**) regulatory gene-gene relations, and after removing each predictor from the ensemble, through leave-one-out cross-validation. For gene-gene relations, the ensemble did not seem to outperform the best tool by itself—ARCHS4’s human expression correlations for positive, and PARMESAN’s indirect predictions for inhibitory relations.

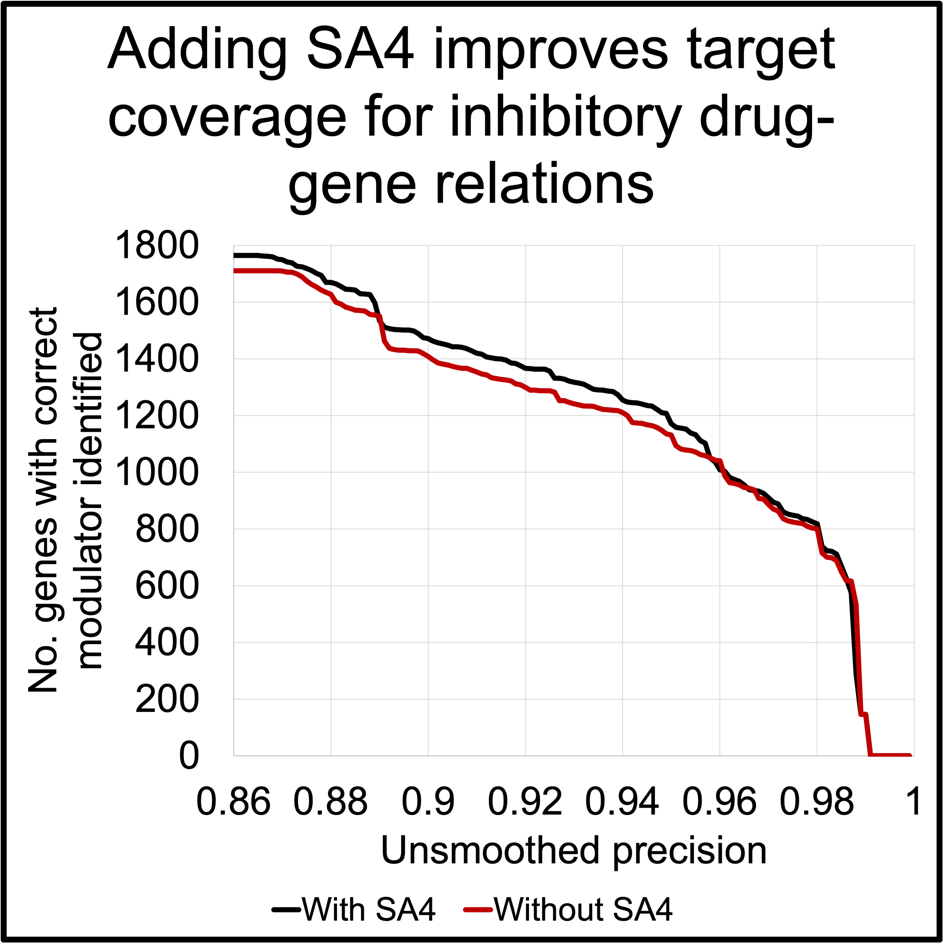

**Supplemental figure 10: Coverage improvement from adding SA4**

We plot the effect that adding SA4 to our repertoire has on the number of targets in DGIdb for which we can correctly identify an inhibitory drug (Y axis), for any unsmoothed precision one is willing to accept (X axis). “With SA4” (black line) uses the same eight drug predictors from the ensemble, and “Without SA4” uses all of them except SA4, demonstrating a clear benefit in coverage upon adding this new predictor.

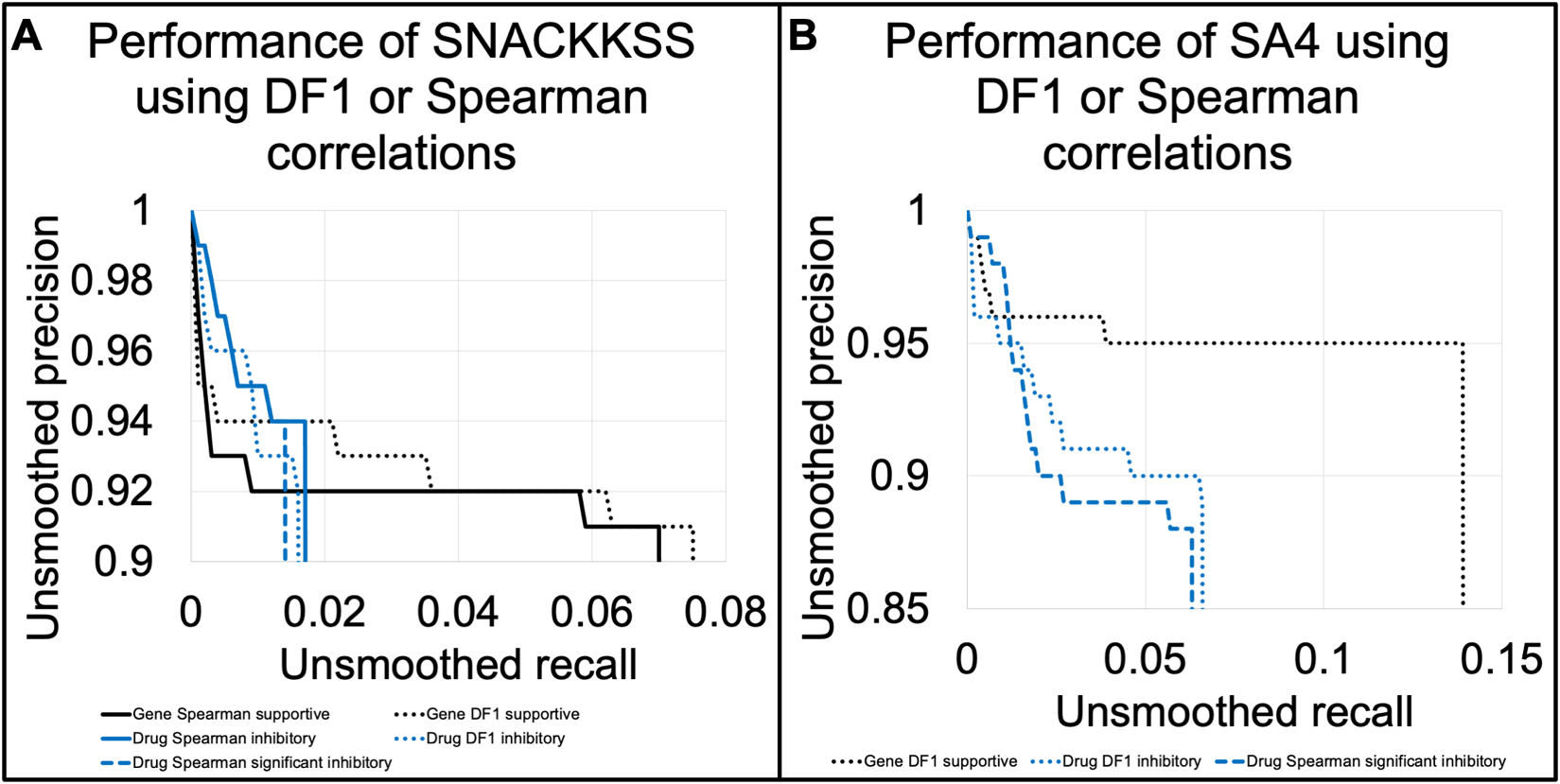

**Supplemental figure 11: Comparison of SNACKKSS’ performance using DF1 or Spearman correlations to match signatures**

We display the unsmoothed precision-recall curves achieved by SNACKKSS (**A**) and SA4 (**B**), using either DF1, a Spearman correlation, or a Spearman correlation requiring a p-value below 0.05 to match signatures. We only plot a predictor if it prioritized correct gold-standard (DGIdb and Reactome) relations over incorrect ones, with an unadjusted log-rank p < 0.05.

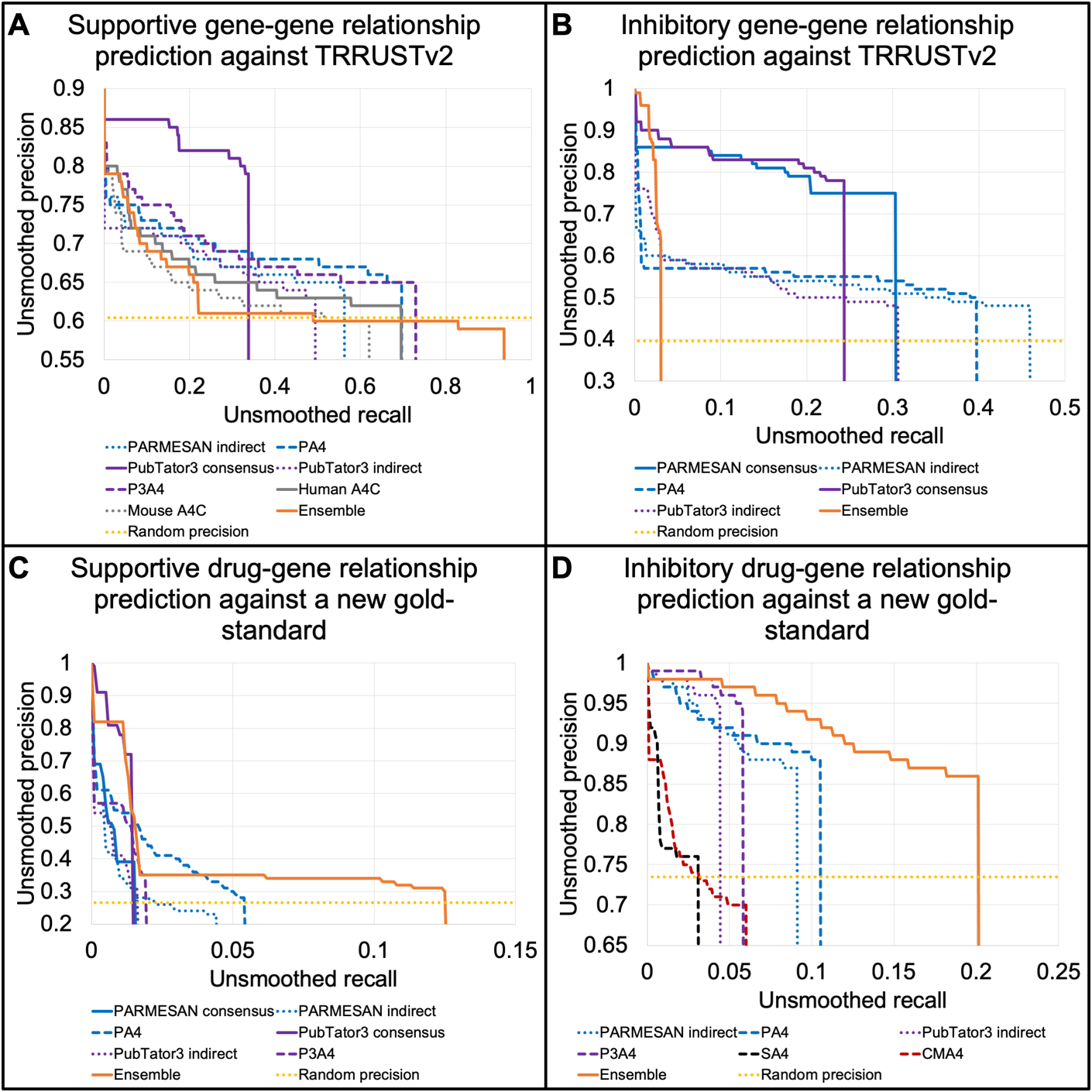

**Supplemental figure 12: Precision and recall of predictive tools against new gold-standard datasets**

We measure the performance (unsmoothed precision and recall) of each predictor in identifying supportive gene-gene (**A**), inhibitory gene-gene (**B**), supportive drug-gene (**C**), and inhibitory drug-gene relations (**D**). The “Ensemble” predictor has each predictive tool estimate its own precision against Reactome (gene-gene) and DGIdb (drug-gene relations), and then believes whichever one has the highest estimated precision in predicting a given relation in the new gold standard described in Supplemental table 20. We only plot a predictor if it prioritized correct relations over incorrect ones, with an uncorrected log-rank p-value below 0.05. We also display the precision that one would achieve through random guessing (“Random precision”, equal to the fraction of the gold-standard dataset that matches the predicted directionality).

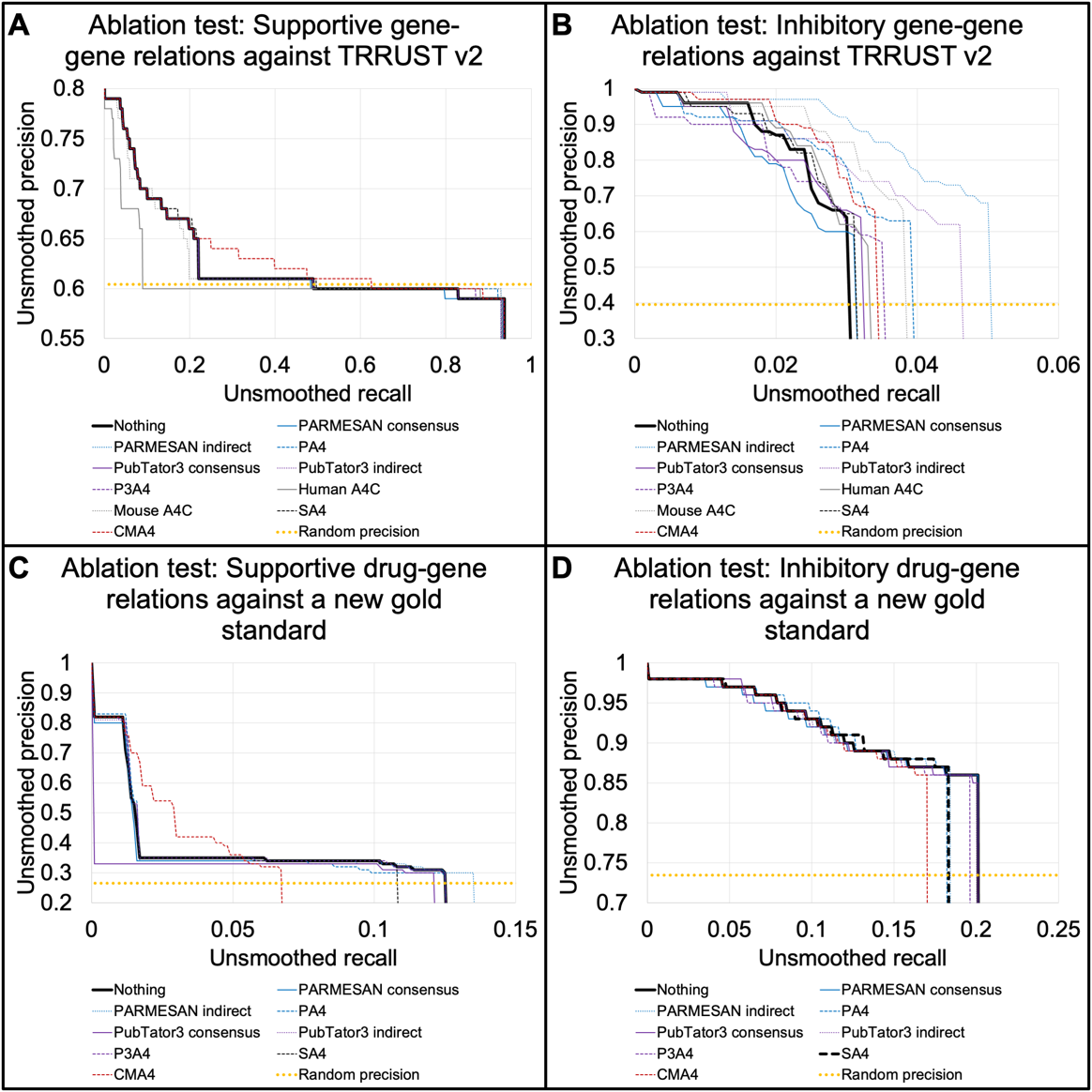

**Supplemental figure 13: Ablation test against new gold-standard datasets**

We run an ablation test, where we plot the unsmoothed precision-recall curve of the ensemble predictor, as is and after removing each predictive tool from consideration. The ensemble estimates each predictor’s accuracy against Reactome (gene-gene) and DGIdb (drug-gene relations) to determine which tool is the most confident, and the precision and recall plotted here are based on a comparison to new gold-standard datasets of gene-gene and drug-gene relations, as described in Supplemental table 20. We separately plot predictions of supportive gene-gene (**A**), inhibitory gene-gene (**B**), supportive drug-gene (**C**), and inhibitory drug-gene (**D**) relations.

3. Hugging Face Token classification. *Transformers*.

4. Hugging Face Text classification. *Transformers*.

5. Deisseroth,C.A., Lee,W.-S., Kim,J., Jeong,H.-H., Dhindsa,R.S., Wang,J., Zoghbi,H.Y. and Liu,Z. (2023) Literature-based predictions of Mendelian disease therapies. *The American Journal of Human Genetics*, 10.1016/j.ajhg.2023.08.018. PMID: 37741276.

6. Wei,C.-H., Allot,A., Lai,P.-T., Leaman,R., Tian,S., Luo,L., Jin,Q., Wang,Z., Chen,Q. and Lu,Z. (2024) PubTator 3.0: an AI-powered literature resource for unlocking biomedical knowledge. *Nucleic Acids Res*, **52**, W540–W546.

7. Wu,G., Feng,X. and Stein,L. (2010) A human functional protein interaction network and its application to cancer data analysis. *Genome Biol*, **11**, R53.

8. Cannon,M., Stevenson,J., Stahl,K., Basu,R., Coffman,A., Kiwala,S., McMichael,J.F., Kuzma,K., Morrissey,D., Cotto,K., *et al.* (2024) DGIdb 5.0: rebuilding the drug-gene interaction database for precision medicine and drug discovery platforms. *Nucleic Acids Res*, **52**, D1227–D1235.
